# Functional divergence between the two cerebral hemispheres contributes to human fluid intelligence

**DOI:** 10.1101/2024.04.05.586081

**Authors:** Xinyu Liang, Junhao Luo, Liyuan Yang, Deniz Vatansever, Elizabeth Jefferies, Gaolang Gong

## Abstract

Hemispheric lateralization is linked to potential cognitive advantages. It is considered a driving force behind the generation of human intelligence. However, establishing quantitative links between the degree of lateralization and intelligence in humans remains elusive. In this study, we propose a framework that utilizes the functional aligned multidimensional representation space derived from hemispheric functional gradients to compute between-hemisphere distances within this space. Applying this framework to a large cohort (N=777 from the Human Connectome Project), we identified high functional divergence across the two hemispheres within the frontoparietal control network. We found that both global divergence between the cerebral hemispheres and regional divergence within the multiple demand network were positively associated with fluid composite score and partially mediated the influence of brain size on individual differences in fluid intelligence. Together, these findings illuminate the profound significance of brain lateralization as a fundamental organizational principle of the human brain, providing direct evidence that hemispheric lateralization supports human fluid intelligence.

## Introduction

The left and right hemispheres of the human brain are not mere duplicates; structural asymmetry and functional lateralization are well demonstrated (*1–4*). While previous studies using multiple techniques, especially functional magnetic resonance imaging (fMRI), have revealed the pivotal role of hemispheric differences in supporting various human cognitive functions (*5*, *6*), the origin of these functional dissociations between the hemispheres remains enigmatic and controversial. A prevailing perspective posits that this divergence is rooted in evolution, and is intimately linked with the development of human intelligence (*7*). This view suggests that to satisfy the varying needs of different tasks in a complex world, the human brain allocates functions asymmetrically across the two hemispheres to optimize processing time (*8–10*). This specialization not only enhances diversity in information processing but also supports our capability for advanced cognition (*11–13*). According to this hypothesis, a certain level of “functional divergence” between the two hemispheres contributes to human intelligence, particularly for more fluid components (*14*). Despite mounting evidence from evolutionary and comparative studies to support the enhanced cognitive capacity by brain lateralization (*15–17*), there is little data in the literature that directly demonstrates a relationship between hemispheric functional divergence and fluid intelligence in humans.

Various hemispheric lateralization patterns in brain organization (*5*, *18*, *19*) have been observed in previous studies based on resting-state functional connectivity (rs-FC), which revealed that the degree of lateralization is associated with specific aspects of cognitive performance, such as language comprehension or visuospatial ability (*5*). These measures capture isolated aspects of functional divergence across the two hemispheres, and as such, they are insufficient for understanding the contribution of hemispheric lateralization to overall intellectual capacity. Moreover, the commonly adopted approach for comparing functional lateralization often involves using flipped or anatomically aligned hemispheres based on landmarks (*5*, *20*). This practice could result in a misalignment of the functional organization across the hemispheres and hamper the accurate estimation of hemispheric functional divergence. Precise estimation of functional distinctions between hemispheres across the entire cortex requires not only structural alignment based on homotopical landmarks but also improved functional correspondence.

To solve these technical issues, we adopted functional alignment (i.e., hyperalignment), aiming at facilitating functional correspondence between the two hemispheres (*21*). In contrast to existing connectivity alignment methods (*22*), we integrated the hemispheric functional connectivity gradients developed in our previous work (*23*). We first estimated functional gradients—i.e., patterns that represent variations in connectivity space that explain the most variance—for the two hemispheres separately, revealing similar but slightly different patterns. This implies that the two hemispheres have a shared functional space, which can then serve as a representational space for functional alignment (*24*, *25*). As a fundamental principle of organization, the cognitive relevance of functional gradients could also facilitate our investigation of fluid intelligence (*26*, *27*).

In summary, we sought to explore the significance of between-hemisphere functional divergence for supporting human fluid intelligence. To address this issue, we used the resting-state fMRI dataset from the Human Connectome Project (HCP) (*28*). We examined within-hemisphere vertexwise resting-state functional connectivity profiles, identified gradients to define a common low-dimensional functional space, and then calculated the between-hemisphere functional distance across homotopic vertices after alignment to this representational space. We tested whether global or vertexwise hemispheric differences contributed to individual differences in fluid intelligence. In addition, previous evidence has shown that brain size plays a crucial role in both general intelligence (*29–31*) and functional lateralization (*32–35*). While larger brains are associated with increased cognitive capacity, a trade-off between efficiency and energy costs constrains brain size (*36*). Hence, we propose that between-hemisphere functional divergence might mediate the influence of brain size on fluid intelligence. Finally, we investigated potential biological factors contributing to hemispheric functional distance.

## Results

### Estimation of functional divergence between left and right hemisphere

To precisely characterize the functional divergence between the two hemispheres, we propose an analytical framework (as shown in Fig. 1) that leverages resting-state fMRI data from 777 right-handed young adults in the HCP cohort (female/male = 428/349; age range = 22-36 years). We identified vertexwise hemispheric functional gradients for each hemisphere on the 10k cortical surface, providing a more detailed and gradual transition than in our previous work (*23*). Specifically, two sets of the top 10 gradients were extracted from individual left and right hemispheres (*37*). All individual hemispheric gradients were subsequently functional aligned through Procrustes rotation to a template representational space derived from the group-level left-right averaged functional connectivity (FC) profiles (Fig. S1), which maintained the internal structure of each hemisphere. The vertexwise between-hemisphere functional distance was measured as the high-dimensional Euclidean distance between each pair of homotopic cortical vertices in the representational space for each participant. The global functional divergence between the left and right hemispheric modules was evaluated by averaging vertexwise between-hemisphere functional distances across the entire cortex. We further performed dimension selection based on the balance between explained variance and test-retest reliability (Fig. S2). We calculated the intraclass correlation coefficient (ICC) of the global metrics and found that it reached a plateau when utilizing the first 6 dimensional axes (global = 0.63, vertexwise = 0.44 ± 0.19). If more dimensions were included, the dimensions with low explanatory rates also introduced more noise components. Hence, a 6-dimensional representation space, capturing 53.47% of the variance in connectivity, was employed to measure between-hemisphere functional distances across individuals. To further mitigate potential noise and reduce comparison time, both vertexwise and global between-hemisphere functional distances were averaged across two sessions for each participant in the subsequent analyses.

**Fig. 1.**
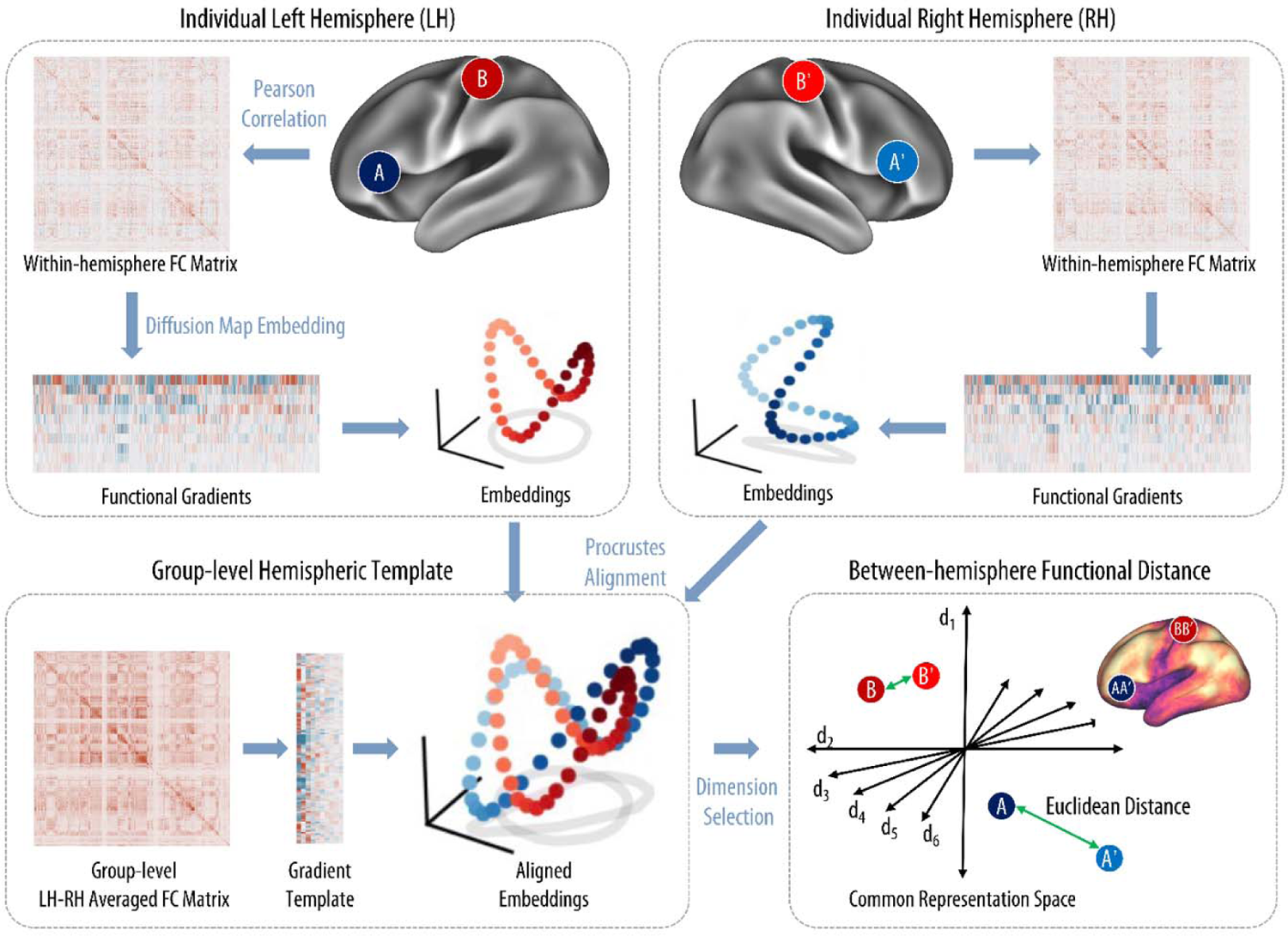
Framework for the computation of between-hemisphere functional distance in a common representational space. To compute between-hemisphere functional divergence, we first calculated functional connectivity (FC) between vertices within each hemisphere based on vertexwise time series data. This yielded hemisphere-specific FC matrices for each hemisphere, resulting in a 9354×9354 matrix after excluding the medial wall. Then we applied the diffusion map embedding method on hemispheric FC matrices to obtain the top 10 functional gradients. These gradients form a low-dimensional representation of the original functional connectivity space, in which the distance represents the functional similarity between vertices. To ensure better comparability between homotopic vertices within the representational spaces of the left and right hemispheres, we employed functional alignment which brought the individual left and right hemisphere embeddings into a group-level common functional space. To determine the optimal dimension for characterizing distance, we used a dimension selection process that balanced explained variance and test-retest reliability. The final between-hemisphere functional divergence was measured as the Euclidean distance between each pair of homotopic vertices in a 6-dimensional common representational space. The sketch plots of embeddings are modified from ref. 80, under the Creative Commons Attribution 4.0 International License (CC BY 4.0).

In addition, we estimated the correlation between the hemispheric differences within each of the top 6 gradients and the between-hemisphere functional distance (Fig. S3). The results indicated that global hemispheric differences in the principal gradient (unimodal to transmodal; *r* = 0.65, *p* < 0.001) and the tertiary gradient (default to task-positive; *r* = 0.68, *p* < 0.001) make particularly large contributions to the individual differences of global functional distance, while the secondary gradient (somatosensory to visual; *r* = 0.32, *p* < 0.001) makes the lowest contribution. The remaining gradients showed medium correlations (above 0.4) with the functional distance. Overall, this analysis suggests that between-hemisphere functional distance is an integration of multiple gradients rather than being solely driven by any single gradient.

### The between-hemisphere functional divergence varies across networks

The spatial distribution of average maps of between-hemisphere functional distance across all participants was first visualized on the cortical surface (Fig. 2a). Notably, the between-hemisphere functional divergence varied widely across the cortex, with greater hemispheric divergence in transmodal regions than in primary and unimodal regions.

**Fig. 2.**
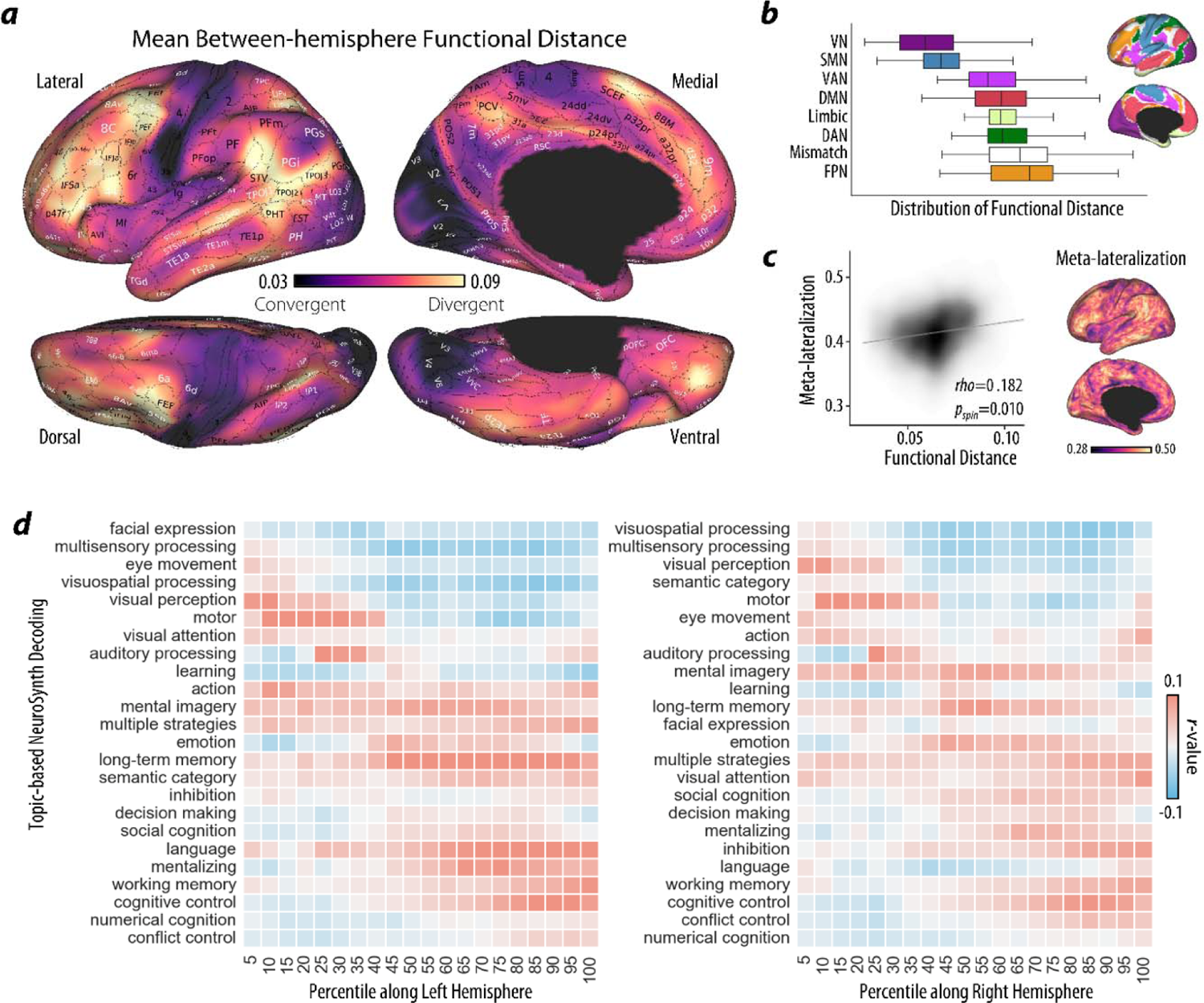
The cortical variation of between-hemisphere functional distance. (**A**). The group average map of between-hemisphere functional distance in the HCP datasets (N = 777). (**B**). We plotted the distribution of between-hemisphere functional distances within each network in a symmetric version of Yeo’s 7 networks. The mismatch zone is defined by mismatch vertices that belong to distinct networks across two hemispheres. (**C**). We calculated the cortical map of the overall functional lateralization index across all 575 cognitive terms. A significant correlation between the cortical map of overall functional lateralization and the between-hemisphere functional distance map was found by a spin test with 10,000 permutations. (**D**). Then we used Neurosynth’s ROI association approach of regions of interest along the between-hemisphere functional distance map with 24 topic terms. The terms were ordered by the positively weighted mean of their location along the left and right hemisphere.

Owing to the asymmetric distribution of Yeo’s 7 networks (*38*), we retained only the homotopic vertices in each network that belong to the same network in both the left and right hemispheres. The networks were arranged according to the degree of hemispheric divergence in descending order from transmodal regions to unimodal regions, including the frontoparietal control network (FPN), dorsal attention network (DAN), default mode network (DMN), limbic network, ventral attention network (VAN), somatomotor network (SMN), and visual network (VN). In addition, vertices belonging to different networks across hemispheres were designated as a mismatch zone. The mismatch zone was situated at the boundaries between adjacent networks and showed a very high between-hemisphere functional distance, second only to that of the FPN. Subsequent analysis indicated that the vertices belonging to the DMN in the left hemisphere and the vertices belonging to the FPN in the right hemisphere contributed the most to this mismatch zone (accounting for 31.76%, Fig. S4).

### The spatial variability of between-hemisphere functional distance reflects hemispheric functional specialization across multiple tasks

We hypothesize that functional divergence between the two hemispheres at rest could serve as a foundational backbone supporting segregated and lateralized processes across a range of tasks. To validate this hypothesis, we first examined whether the spatial distribution of between-hemisphere functional distance aligns with the cortical pattern of overall functional lateralization observed in meta-analytic task activations. We obtained an overall functional lateralization index for the cortical surface across multiple cognitive domains (all 575 cognitive terms) from previous studies based on the Neurosynth database (*2*, *39*) (Fig. 2c). The significance of the spatial correspondence for the between-hemisphere functional distance derived from intrinsic connectivity and the overall functional lateralization index derived from meta-analytic task data was assessed using a spin permutation test, which generates the significance level (denoted as *p*_spin_) based on a null distribution of randomly rotated brain maps that preserve the spatial covariance structure of the original data (using 10,000 permutations). Between-hemisphere functional distance was significantly correlated with the distribution of overall functional lateralization (*rho* = 0.182, *p*_spin_ = 0.010; Fig. 2c).

Furthermore, we conducted a topic term-based meta-analysis along the between-hemisphere functional distance map within 5-percentile bins for each hemisphere (*40*, *41*). The resultant topic terms were sorted by their positively weighted average positions along each hemisphere, revealing a systematic shift from perception to higher-level cognition, especially cognitive control, language, number processing and working memory (Fig. 2d). Although the order of topic terms was relatively consistent across the left and right hemispheres, functions such as language, long-term memory, mentalizing and semantic categorization were found to be closer to the highest level in the left hemisphere than in the right hemisphere, whereas functions such as facial expression discrimination, visual attention and inhibition demonstrated the opposite pattern (Fig. S5).

### Demographic and physiological associations with global between-hemisphere functional distance

At the individual level, we initially focused on the relationships between global between-hemisphere functional distance and demographic and physiological factors, such as age, sex, handedness and brain size, which have been associated with hemispheric functional lateralization in previous studies. Although we regressed out the parameters of in-scanner head motion during preprocessing, we still included the mean framewise displacement as a covariate in all subsequent analyses due to evidence that head motion is a confound of intrinsic connectivity metrics as well as fluid intelligence (*42*). For each participant, the total brain volume was calculated by summing the volumes of gray matter and white matter. For each factor, we employed partial correlation, with the other factors serving as covariates. Although the age range (22–36 years) of the adult participants was relatively narrow, the global functional distance between the two hemispheres was significantly positively correlated with age (*r* = 0.159, *p* < 0.001; Fig. S6a). In terms of sex, the global between-hemisphere functional distance in males was significantly greater than that in females (*F*(_1,749_) = 5.55, *p* = 0.019; Fig. S6b). Global between-hemisphere functional distance was also positively correlated with total brain volume (*r* = 0.238, *p* < 0.001; Fig. 3a), as larger brains showed a greater functional distance between the two hemispheres. There was no correlation between the global between-hemisphere functional distance and handedness score (*r* = 0.052, *p* = 0.155; Fig. S6c).

**Fig. 3.**
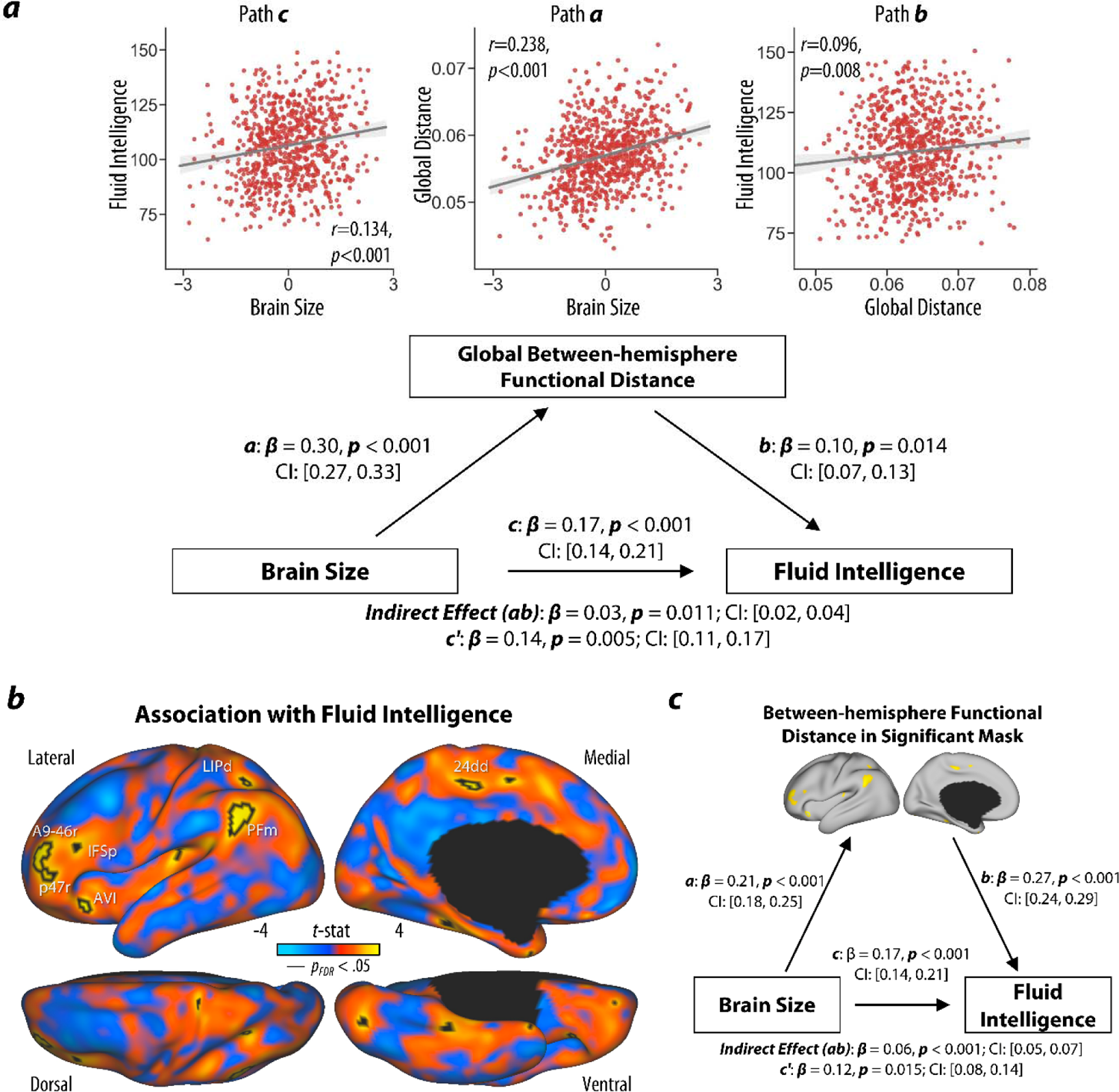
Individual differences in between-hemisphere functional distance mediate the effect of brain size on fluid intelligence. (**A**). We observed significant correlations between brain size, global between-hemisphere functional distance and fluid intelligence after controlling for age, sex, handedness and mean framewise displacement. We tested the significance of the hypothetical mediation pathway, in which global between-hemisphere functional distance could partially mediate the impact of brain size on fluid intelligence. Significance was tested by bootstrapping (10,000 replacements). (**B**). A GLM analysis was performed on vertexwise between-hemisphere functional distances to identify regions associated with intelligence, in which age, sex, handedness, brain size and mean FD were included as covariates. The resultant t maps for fluid intelligence are shown. The regions with gray outlines were significant vertices after FDR correction (one-tailed *p*_FDR_ < 0.05). (**C**). The mean between-hemisphere functional distances within a mask consists of all significant vertices also exhibited a significant mediating effect on the relationship between brain size and fluid intelligence.

### The global between-hemisphere functional distance mediates the effect of brain size on fluid intelligence

We hypothesized that the overall functional divergence between the two cerebral hemispheres positively contributes to enhancing fluid cognitive capabilities. To verify this hypothesis, we explored the association between global between-hemisphere functional distance and fluid intelligence. Here, we utilized the composite score of fluid intelligence from the National Institutes of Health (NIH) Toolbox Cognition Battery (*43*), which combines scores on cognitive flexibility, executive inhibition, episodic memory, working memory, and processing speed. The global between-hemisphere functional distance was positively correlated with the fluid cognition composite score (*r* = 0.125, *p* < 0.001), indicating that participants with a greater hemispheric functional distance had slightly greater fluid intelligence. Given the significant relationship between total brain volume and fluid cognition (*r* = 0.134, *p* < 0.001), we additionally included brain size as a covariate. After controlling for total brain volume, the correlation between global between-hemisphere functional distance and fluid cognition was weaker but still significant (*r* = 0.096, *p* = 0.008). To further explore which cognitive domain makes the strongest contribution to this relationship, we conducted post hoc correlation analyses. Among the five subdomain scores, we only observed a significant positive correlation between cognitive flexibility and global between-hemisphere functional distance (*r* = 0.134 *p* < 0.001; Table S1).

To investigate whether hemispheric difference mediates the influence of brain size on fluid intelligence, we tested a mediation model including total brain volume as the predictor, global between-hemisphere functional distance as the mediator, and the fluid cognition composite score as the outcome (path c, β = 0.17, *p* < 0.001; path a, β = 0.30, *p* < 0.001; path b, β = 0.10, *p* = 0.014). Bootstrap simulation analysis (10,000 times) confirmed a significant indirect effect (a × b = 0.03, 95% confidence interval = [0.02, 0.04], *p* < 0.001; Fig. 3a). Our findings suggest that between-hemisphere functional divergence can partially explain the association between brain size and fluid cognitive ability.

### The role of vertexwise between-hemisphere functional distance in fluid intelligence

As we identified a significant association between global between-hemisphere functional distance and the fluid composite score, we further aimed to investigate which specific brain regions are implicated in this relationship. We employed a general linear model (GLM) capturing between-hemisphere functional distance at the vertexwise level. After controlling for age, sex, handedness, head motion and brain size, we found that better fluid intelligence was associated with between-hemisphere functional distance in the inferior parietal cortex (area PFm), anterior dorsolateral frontal cortex (area a9-46v), orbital prefrontal cortex (area p47r), anterior ventral insula (area AVI), dorsal part of the lateral intraparietal region (area LIPd), and dorsal part of the ventral anterior cingulate (area 24dd) according to multimodal parcellation (*44*) (Fig. 3b). These regions were mainly located in the FPN or mismatch zone in Yeo’s 7 networks (Fig. S7). For instance, the vertices comprising significant homotopic pairs in PFm were affiliated with distinct networks, as the left component was in the DMN, while the right component was in the FPN.

In addition, we conducted a post hoc correlation analysis on all significant vertices (179 vertices) as a combined mask. The average between-hemisphere functional distance in the combined mask exhibited a positive association with the fluid cognition composite score (*r* = 0.265, *p* < 0.001). To test whether the average between-hemisphere functional distance in this significant mask also mediates the effect of brain size on fluid intelligence, we performed a similar mediation analysis as above. Our findings demonstrated a significant partial mediating effect of between-hemisphere functional distance on the relationship between brain size and fluid intelligence (path c, β = 0.17, *p* < 0.001; path a, β = 0.21, *p* < 0.001; path b, β = 0.27, *p* < 0.001; a × b = 0.06, 95% confidence interval = [0.05, 0.07], *p* < 0.001; Fig. 3c).

### Potential biological determinants affecting between-hemisphere functional distance

We evaluated how strongly between-hemisphere functional distance was associated with cortical microstructure reflecting myelination. The cortical T1w/T2w map (Fig. 4a) has been proposed as an in vivo measure that is sensitive to regional variation in gray matter myelin content (*45*). We found that cortical between-hemisphere functional distance was significantly correlated with group-averaged T1w/T2w (averaged across the left and right sides, *rho* = −0.55, *p*_spin_ < 0.001), indicating a potential contribution of myelination to between-hemisphere distance: the most functionally divergent regions across the left and right hemisphere are also the least myelinated. In addition, we tested the hypothesis that between-hemisphere functional divergence is driven by evolutionary change. Our results indicated that the spatial pattern of between-hemisphere functional distance is well aligned with evolutionary cortical expansion indices (*rho* = 0.56, *p*_spin_ = 0.001), which reflect cortical expansion in humans relative to macaques (Fig. 4b) (*46*). Moreover, we found no significant correlation between between-hemisphere functional distance and developmental expansion (*rho* = 0.30, *p*_spin_ = 0.035), which is a metric that reflects cortical expansion in human adults relative to human infants (Fig. 4c). Together, our results demonstrate that between-hemisphere functional divergence could be an evolutionary outcome suggesting that it may confer functional advantages that benefit survival and reproduction.

**Fig. 4.**
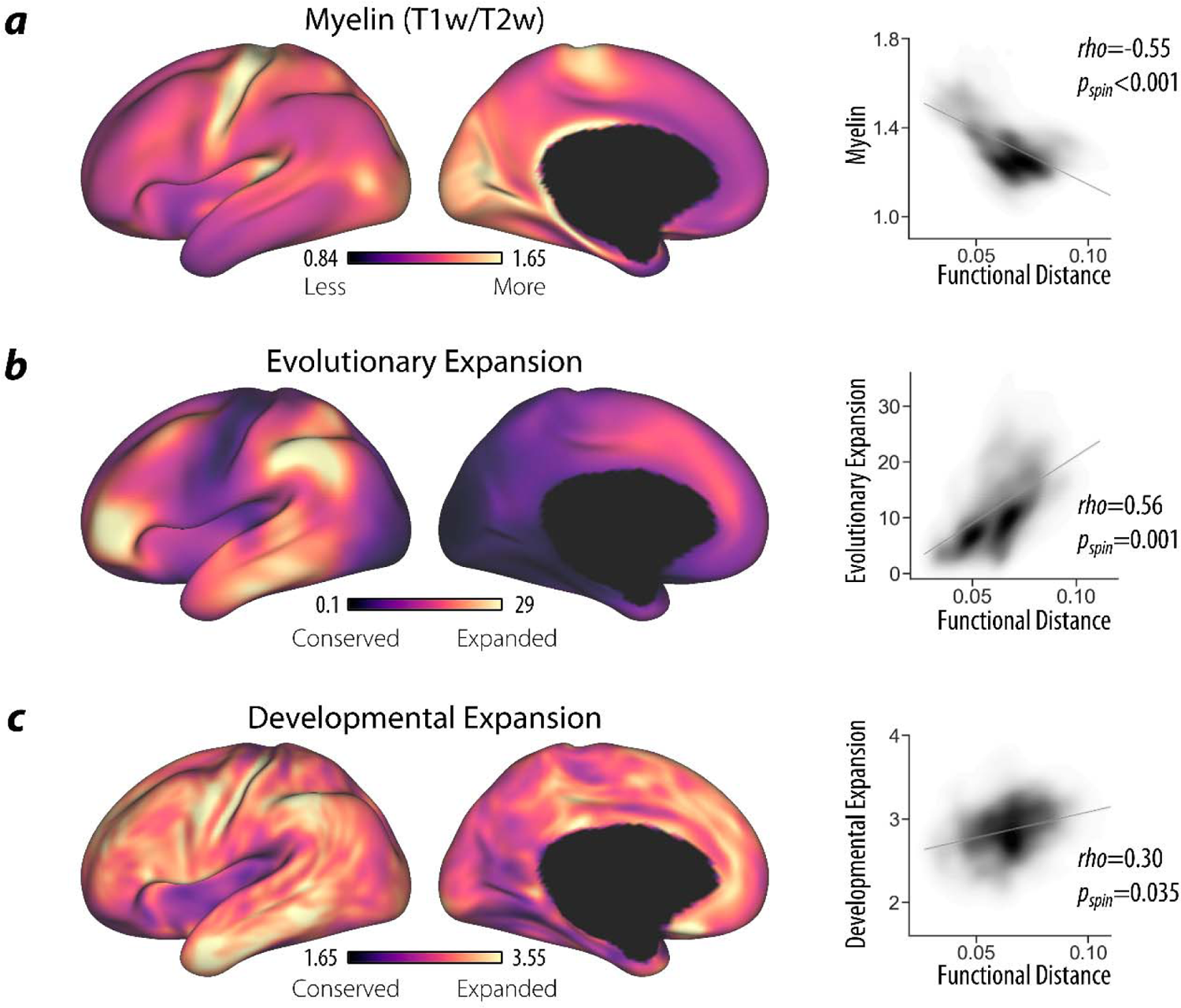
Spatial variability in between-hemisphere functional distance relates to biological maps. Cortical maps of myelin (**A**), evolutionary expansion (**B**), and developmental expansion (**C**) were utilized to test the potential biological basis for between-hemisphere functional distance. Their spatial correspondences with group-averaged between-hemisphere functional distances are plotted. The significance level was determined by a spin test with 10,000 permutations with the Bonferroni correction (*p*_spin_ < 0.0125).

### Validation analyses

To assess the robustness of our findings, we repeated the main analyses while regressing out the global signal of the whole brain (Fig. S8 and Table S2) and our key results were replicated. In addition, to estimate the stability of the association between global between-hemisphere functional distance and fluid intelligence and how this is influenced by sample size (*47*), we examined 1000 random subsamples at a range of sampling rates (15 intervals evenly spaced from 50 to 750). The association becomes stable once the sample size reaches 650 (Fig. S9). To validate a general vertexwise association between between-hemisphere functional distance and fluid intelligence, we adopted a leave-one-subject-out cross-validation (LOOCV) analysis. The observed fluid intelligence score and predicted scores generated by support vector regression showed significant positive correlation (*r* = 0.146, *p* < 0.001, Fig. S10).

To substantiate the efficacy of our proposed framework, we also considered whether alternative measures of hemispheric functional difference would identify relationships with fluid intelligence. Specifically, we computed the homotopic FC (*48*) and the lateralization of within-hemisphere FC strength (*18*) for each pair of homotopic vertices. Homotopic FC describes the functional synchrony of time series data, while the lateralization of within-hemisphere FC strength reflects the absolute degree of hemispheric differences in functional integration within each hemisphere. Compared to between-hemisphere functional distance, homotopic FC displayed a comparable ICC across scan sessions (global = 0.68, vertexwise = 0.60 ± 0.14), whereas the ICC of lateralization of within-hemisphere FC strength was much lower (global = 0.52, vertexwise = 0.18 ± 0.07, Fig. S11). Notably, neither global metric demonstrated statistically significant correlations with the fluid cognitive composite (homotopic FC: *r* = −0.021, *p* = 0.569; lateralization of within-hemisphere FC strength: *r* = 0.007, *p* = 0.854; Table S3). This demonstrates the value of our approach based on dimensions of intrinsic connectivity; by focusing on a small number of variables that explain the most functional variance, it is possible to discern hemispheric differences that predict individual differences in cognition.

## Discussion

By leveraging state-of-the-art functional alignment and connectivity gradient techniques, we introduced an analytical framework to assess between-hemisphere distance in a functional representational space, which effectively quantified the level of functional divergence in each brain region and showed spatial alignment with lateralized response patterns in task meta-analysis. Our findings revealed a greater functional distance between the two hemispheres within the frontoparietal network and between mismatched pairs of vertices belonging to the left DMN and right FPN. Notably, these regions with high between-hemisphere functional distances are involved in higher-order cognitive functions, particularly cognitive control. We directly verified the association between global between-hemisphere functional distance and fluid intelligence, thereby providing the empirical evidence for the hypothesis that hemispheric specialization affords functional advantages across multiple cognitive domains in humans. Global and regional between-hemisphere functional distance also mediated the influence of brain size on fluid intelligence. Finally, we demonstrated the contribution of structural features to the spatial pattern of between-hemisphere functional distance, including the effects of cortical myelination and evolutionary area expansion. These discoveries collectively bridge a significant gap, elucidating the intricate mechanisms behind the formation of functional hemispheric lateralization to support fluid cognitive intelligence.

The cortical pattern of between-hemisphere functional divergence shows high similarity with previous hemispheric lateralization patterns observed across multiple levels of functional activity and connectivity, including meta-analytic activation (*2*), within-hemisphere integration (*18*, *49–51*), between-hemisphere integration and segregation (*5*, *6*, *19*, *52*) and hemispheric gradients (*23–25*). However, our results also provide a more holistic delineation of the complex structure of hemispheric lateralization by simultaneously considering multiple gradients that reflect the core dimensions of functional representational space; in this way, we are able to better identify the impact of lateralization on fluid intelligence. In comparison to our previous work (*23*), this high-dimensional space captures subtle interactions among gradients and largely avoids the methodological problems of traditional asymmetry indices ([L-R]/[L+R]), such as arbitrary cut-offs that classify dominance, non-normal distributions, and inapplicability of negative values (*53*, *54*). The proposed functional distance provides more accurate, sensitive and reliable estimates of brain lateralization, which can be applied in many contexts that examine functional hemispheric dissociations.

Leveraging this metric of between-hemisphere functional distance, we revealed an association between hemispheric lateralization and individual differences in fluid intelligence. This link is consistent with, and helps to unify, previous findings suggesting that specific cognitive processes are supported by lateralized patterns of functional organization, including visuospatial attention and long-term memory (*5*, *13*, *49*). These benefits to specific aspects of cognition might facilitate fluid abilities. In the two-component theory of intelligence (*14*), fluid intelligence is thought to be associated with metacognition and the executive control of cognitive processing to achieve adaptive cognition (*55*). Our vertexwise analysis revealed that lateralization in the rostrolateral prefrontal cortex (RLPFC) and the anterior extent of the inferior parietal lobule (IPL) are significantly related to fluid intelligence. These regions fall within the FPN, which is associated with the integration of information across networks to facilitate flexible and adaptive cognition (*56*). Studies have shown that bipartite subsystems of the FPN differentially support distinct aspects of cognitive control (*24*, *56*, *57*). The Control-A subsystem (including the dorsolateral prefrontal cortex) is more closely coupled with attention regions in the DAN, while the Control-B subsystem (including the IPL and the RLPFC) is more closely coupled with memory regions located in the DMN. Adaptive cognition needs to integrate controlled retrieval of meaningful information from memory with external goal-driven attention. Depending on the demands of cognitive tasks, these subsystems within the FPN may induce distinct patterns of lateralization. For example, the control of semantic cognition as well as internal states could involve a more leftward asymmetric pattern of network coupling than the control of spatial attention (23). We speculate that these segregated patterns of connectivity for control subsystems across hemispheres could give rise to functional benefits such as reduced interference and shorter network restructuring time.

Our findings extend key theoretical accounts of human fluid intelligence. The identified regions, including RLPFC and IPL, are largely consistent with the theoretical networks that underpin human intelligence. The parieto-frontal integration theory (P-FIT) proposed by Jung and Haier (*58*), suggests that a lateralized distributed network, including the dorsolateral prefrontal cortex and superior and inferior parietal lobule, integrates knowledge to support reasoning and problem solving. Similarly, the multiple demand (MD) system located in the prefrontal and parietal cortex has been suggested to be a key network supporting fluid intelligence (*59*). The MD system is typically assumed to be bilateral due to the methodology used to identify this network, which relies on shared activation patterns across a wide range of tasks (*60*); these tasks may manifest varying degrees of lateralization but become symmetrized after averaging across different activation patterns. A latest network neuroscience theory also suggests that the dynamics and flexibility of frontoparietal network are strongly associated with the fluid intelligence (*61*). Our discoveries extend these theories by showing that frontoparietal system can be asymmetrically active and that between-hemisphere functional divergence within this system acts to support fluid intelligence.

The positive association of brain size with general intelligence has been validated in multiple in vivo human studies, including meta-analyses (r = 0.24, N = 8,036 from 148 independent samples) (*30*), and in large cohorts including the HCP (r = 0.24, N = 896) (*31*) and the UK Biobank (r = 0.28, N = 7,201)(*29*). Despite the robustness of this association, the reasons why and how brain size could affect human intelligence are still debated (*30*). Brain size could be a proxy for neuron number, which may have contributed to the evolution of higher cognitive abilities in primates. However, given species-specific constraints (reflecting the ratio of brain-body size), neural capacity could be enhanced through left versus right functional differences (*9*, *62*, *63*). Many studies have shown a relationship between whole-brain size and brain lateralization (*32*, *33*, *64*). Our findings provide vital evidence to support this hypothesis in human, suggesting that divergent functions across hemispheres are beneficial for human fluid intelligence, especially for people who have larger brains. This may be a consequence of both a reduction in the time-consuming transmission of information and the prevention of interference between the hemispheres, maximizing the efficiency of neural processing.

Our findings also revealed that prominent between-hemisphere divergence is located in regions linked to dramatic cortical expansion during evolution (*65*). Evolutionary cortical expansion has been implicated in human cognitive capacities, such as language (*66*). Our results further suggest that evolutionary cortical expansion, rather than developmental processes, plays a considerable role in shaping the spatial distribution of functional lateralization across the entire cortex. The significant association between cortical myelination and between-hemisphere divergence is also compatible with the idea that more lateralized regions have a smaller density of interhemispheric connections (*2*, *67*).

There are, of course, potential limitations of this study. First, extracting vertexwise gradients has high computational cost, and future work is required to optimize the balance between spatial resolution and computational cost to make this analysis approach more efficient. Second, structural white matter connections are likely to be relevant to between-hemisphere functional distance, and further research is needed to explore how structural and functional connectivity patterns contribute to functional divergence. Third, we confined our analyses to healthy young right-handed adults and further research is needed to generalize our conclusions to left-/bilateral-handed adults, children or elderly individuals. Finally, the lifespan trajectory of between-hemisphere functional distance warrants future investigation to reveal its development.

In sum, this study highlights the significant contribution of between-hemisphere functional lateralization to human intelligence, and the method we developed could serve as a catalyst for the continued exploration of the origins and significance of brain lateralization.

## Materials and Methods

### Dataset

The participants in the present study were from the HCP S1200 release (*28*). The HCP was reviewed and approved by the Institutional Ethics Committee of Washington University in St. Louis, Missouri. All participants gave written informed consent. For more details, please refer to Van Essen et al. 2013. In our study, we excluded 146 HCP participants with quality control issues (including codes A, B and C from the HCP minimal preprocessing pipeline). A further 190 HCP participants were excluded owing to their lack of rs-fMRI data. To avoid the possible confounding effect of handedness, we further excluded 18 HCP participants with bilateral handedness and 75 HCP participants with left handedness (setting the scores on the Edinburgh Handedness Questionnaire of ± 20 as the threshold). The final sample included 777 right-handed young adults (female/male = 428/349; age range = 22-36 years; handedness [mean ± SD] = 79.55 ± 17.85).

### Cognitive assessment

Participants in the HCP dataset were subjected to a wide range of validated cognitive tests derived from the NIH Toolbox Cognition Battery (NIH-TCB, http://www.nihtoolbox.org), which mainly focused on five domains: language, executive function, episodic memory, processing speed, and working memory. Based on the specific tests above, three composite scores were derived from the task scores: crystallized cognition composite, fluid cognition composite, and total cognition composite (*43*). Here, we focused only on the fluid cognition composite, which broadly assesses processing speed, memory, and executive functioning and has demonstrated remarkable reliability and robust construct validity (*68*). It comprises scores on the Dimensional Change Card Sort for executive cognitive flexibility, the Flanker task for executive inhibition, the picture sequence memory task for episodic memory, the list sorting task for working memory, and the Salthouse pattern comparison task for processing speed. A detailed description of each cognitive test and its scoring can be found on the HCP website (http://www.humanconnectome.org/documentation/Q1/behavioral-measuresdetails.html).

### MRI acquisition

All MRI data were collected using the same 3T Siemens Skyra magnetic resonance machine at the University of Washington, with a 32-channel head coil (*28*). Specifically, rs-fMRI was acquired by using a gradient-echo echo planar imaging (GE-EPI) sequence with the following parameters: repetition time (TR) = 720 ms, echo time (TE) = 33.1 ms, flip angle (FA) = 52°, bandwidth = 2290 Hz/pixel, field of view (FOV) = 208 × 180 mm^2^, matrix = 104 × 90, voxel size = 2 × 2× 2 mm^3^, multiband accelerated factor = 8, slices = 72, and total scan time of 1200 frames = 14 min and 24 s. During the scan, participants were asked to open their eyes and stare at a white cross on the screen with a black background. There were two rs-fMRI sessions (REST1 and REST2) acquired on the two consecutive days, each including the two runs with a left-to-right (LR) and a right-to-left (RL) phase encoding direction. The T1-weighted images were acquired by using a magnetization-prepared rapid gradient-echo imaging (MPRAGE) sequence with the following parameters: TR = 2400 ms, TE = 2.14 ms, reversal time (TI) = 1000 ms, FA = 8°, FOV = 224 × 224 mm^2^, voxel size 0.7 mm isotropic, and total scan time = 7 min and 40 s. The T2-weighted images were acquired by using a T2-SPACE sequence with the following parameters: TR = 3200 ms, TE = 565 ms, FA = 8°, FOV = 224 × 224 mm^2^, voxel size 0.7 mm isotropic, and total scan time = 8 min and 24 s.

### Resting-state fMRI postprocessing

We used the HCP minimally preprocessed data (*69*). For rs-fMRI, the procedures of the HCP minimal preprocessing pipeline (version 2.0) include magnetic gradient distortion correction, EPI distortion correction, nonbrain tissue removal, Montreal Neurological Institute (MNI) standard space registration, and intensity normalization. The resulting data were further denoised by using independent component analysis (ICA) with the FIX tool (*70*, *71*), which can effectively distinguish and remove the components of spatiotemporal signals caused by nonneuronal or structural noise, especially head movement. The individual resulting time series were projected to the standard 32k_fs_LR surfaces, on which the homotopic vertices are topographically corresponding.

We subsequently implemented several postprocedures on the minimally preprocessed rs-fMRI data. First, linear detrending was performed to minimize the effects of low-frequency drift (*73*). Second, a set of nuisance variables that were not related to neural signals, including average white matter and cerebrospinal fluid signals, was regressed out. Given the controversy regarding the removal of global signals (*74*), we did not regress out the global signals in our main results. Third, we performed temporal bandpass filtering (0.01-0.1 Hz) on the time series to minimize high-frequency physiological noise (*75*). To better remove motion artifacts, we further performed data censoring (i.e., scrubbing) without interpolation (*76*). For each participant, we censored the time frames whose framewise displacements (FDs) exceeded 0.5 mm and the frames one before and two after. Finally, the residual BOLD time series on the 32k_fs_LR surface were downsampled to a homemade 10k_fs_LR surface to reduce computing resource consumption. Following the steps described in the manual for surface resampling (https://wiki.humanconnectome.org/download/attachments/63078513/Resampling-FreeSurfer-HCP_5_8.pdf), we created an identity sphere with 10242 vertices to maintain the hemispheric correspondences between homotopic vertices. The ADAP_BARY_AREA method was used for the metrics to reduce the errors caused by resampling.

### Functional gradients estimation and between-hemisphere functional alignment

Using preprocessed rs-fMRI images, we estimated hemispheric functional gradients following our previous work (*23*). Rather than focusing on the region level, we used the vertexwise connectivity in this study to obtain a high-resolution delineation. After masking out the combined medial wall across the two hemispheres, the entire cortical surface included 9354 paired homotopic vertices. To remove the influence of imaging phase encoding in each rs-fMRI session, we concentrated the time series of the LR and RL phase encoding runs (*77*). For each pair of vertices, the Pearson correlation coefficient (converted to Fisher’s *Z* values) of the time series was computed as the FC strength. Thus, for each subject, two 9354×9354 within-hemisphere FC matrices (left and right) were generated. For each vertex, the within-hemisphere FC profile was defined as a vector (i.e., the corresponding column from the within-hemisphere FC matrix above), representing its connectivity to other vertices within the same hemisphere.

For each hemisphere, we utilized the BrainSpace toolbox to estimate functional gradients (*37*). First, the within-hemisphere FC profile vector of each region was thresholded by retaining the 10% of connections with greatest strength and setting the remaining connections to zero, as was done previously (*40*). For each pair of vertices within the same hemisphere, the normalized angle similarity coefficient was calculated to quantify the similarity of their thresholded FC profile vectors (*78*). The similarity matrix was subsequently subjected to the diffusion map embedding algorithm, which yielded multiple continuous components, i.e., functional gradients (*79*). Here, the parameter α of manifold learning was set to 0.5 (*40*). The resultant functional gradients can be seen as a collection of continuous axes as a low-dimensional representation of the initial high-dimensional connectivity space. For each vertex, its scores on specific gradient components along those continuous axes formed a set of coordinates depicting its location in the low-dimensional representation space.

To ensure the comparability of the representational spaces between the left and right hemispheres, as well as across participants, we generated a group-level hemispheric gradient template for subsequent functional alignment (*21*). Specifically, we averaged all left and right within-hemisphere FC matrices across all participants and then generated template gradients from this average within-hemisphere FC matrix. We retained the first 10 gradient axes as the template space, accounting for the majority of the variance (approximately 66.15% in REST1 and 65.85% in REST2) in the initial connectivity space. Procrustes rotation was utilized to align the first 10 gradient axes for both the left and right hemispheres of each individual with the group-level template space while keeping the internal structure of each hemisphere unchanged (*80*).

### Between-hemisphere functional distance estimation

Inspired by a previous study (*81*), the between-hemisphere functional distance was measured as the Euclidean distance between each pair of homotopic vertices in the common representation space. For each participant, the global functional distance between the left and right hemispheres was further computed by averaging the vertexwise between-hemisphere functional distances across the entire cortex, which is equivalent to the Euclidean distance between the hemispheric centroids. To determine the optimal dimension for characterizing the distance, we proceeded with a dimension selection process to balance the explained variance and the test-retest reliability. We calculated the intraclass correlation coefficient (ICC) of the global between-hemisphere functional distance between the two scan sessions and found that it had already reached a plateau when utilizing the first 6 dimensional axes. If more dimensions had been included in the calculation, the dimensions with low explanatory rates could have introduced more noise components. Finally, we used the between-hemisphere functional distances calculated in the 6-dimensional functional space (as shown Fig. S1), which accounted for approximately 53.47% of the total variance in REST1 and 52.49% of the total variance in REST2. The between-hemisphere functional distances were further averaged across two sessions for each participant to reduce potential noise as well as comparison time.

To estimate the contribution of hemispheric differences along each gradient dimension to the final high-dimensional distance, we calculated the similarity between the proposed between-hemisphere functional distance and the absolute hemispheric difference in score for each of the first 6 gradients. The similarity was calculated by Pearson correlation for both the spatial distribution within each participant and the global measurement across participants.

### Hemispheric consistency of Yeo’s 7 networks

The original Yeo’s 7 networks (the somatomotor, visual, dorsal attention, ventral attention, limbic, and frontoparietal networks, and the default mode networks) exhibited an asymmetrical distribution. The atlas of Yeo’s 7 networks was initially resampled to the homemade 10k_fs_LR surface. For each network, we preserved the homotopic vertices whose identity labels belonged to that specific network. In cases where there were vertices with mismatched identities belonging to different networks in the two hemispheres, we designated those vertices as a mismatch zone. The mismatch zone situated along the boundaries between adjacent networks and the DMN in the left hemisphere and the FPN in the right hemisphere contributed the most to the observed zone (accounting for 31.76%). We plotted the constituents of the mismatch zone (Fig. S4) by using Circos (http://mkweb.bcgsc.ca/tableviewer/).

### Neurosynth decoding of the between-hemisphere functional distance map

To determine the cognitive functions associated with the investigated between-hemisphere functional distance map, we conducted two distinct types of meta-analyses utilizing the Neurosynth database (http://Neurosynth.org). Following the methods of a recent study, we estimated the overall functional lateralization index across cognitive domains (*2*). Initially, we selected available cognitive terms (575 in total) from the current Neurosynth database. We applied the Neurosynth tool to generate a whole-brain meta-activation image for each cognitive term in MNI space. After aligning these meta-activation images to a symmetric image template, we projected the values onto the standard 32k_fs_LR surface and then resampled them to the 10k_fs_LR surface. Only pairs of cortical homotopic vertices exhibiting positive meta-activation were retained for each term. Then the functional lateralization indices of the meta-activation values were calculated by using the conventional lateralization index formula (LI = abs[L – R]/[L + R]). To generate an overall functional lateralization map spanning all cognitive domains, we simply averaged the nonzero values across all 575 cognitive terms for each vertex. The resulting map was subsequently utilized to assess the spatial correspondence between brain maps.

In addition, we performed meta-analytic functional decoding according to the method of Margulies et al. (*40*). The group-averaged cortical between-hemisphere functional distance map was divided into increments of five percentiles. From these 20 maps, spanning 0% to 5% up to 95% to 100%, we generated region of interest (ROI) masks.

These masks were projected onto the volumetric MNI152 standard space based on the left and right surfaces, and the resulting volumetric maps were converted into binary forms for input into the meta-analysis. The feature terms used in this study were derived from the 50 sets of topic terms (*82*), and 24 topic terms related to cognitive functions were manually selected following the method of a previous study (*40*). For each ROI map, the output of the analysis yielded a correlation linked to the selected feature term. In each hemisphere, these terms were subsequently arranged based on their positively weighted means for visualization purposes. We additionally compared the hemispheric differences (left-right) between the correlations of each topic term. We visualized the terms according to their average differences in correlation in ascending order (Fig. S5). The meta-analysis was implemented through Neurosynth’s ROI association approach in NiMARE (v0.0.11).

### Estimation of individual brain size

The total brain volume was defined as the estimated brain size, which was calculated as the sum of the total amount of intracranial brain tissue (gray and white matter). The FreeSurfer (version 5.3) pipeline in HCP minimal processing had already performed skull stripping and tissue segmentation (*69*). We used the data from the ‘FS_IntraCranial_Vol’ column in the sheets provided by the HCP.

### Cortical maps for microstructure and expansion

Group-averaged T1w/T2w maps (N = 1096) were obtained from the HCP. The hemispheric T1w/T2w maps were further averaged across the left and right hemispheres, resulting in a single map representing the myelin information for each vertex (Fig. 4a)(*45*). Cortical expansion maps were estimated by Hill and colleagues (*46*). The evolutionary expansion index was evaluated in humans relative to macaques (Fig. 4b), and the developmental expansion index (Fig. 4c) was estimated in human adults relative to human infants. These cortical maps were resampled onto our homemade 10k_fs_LR surface.

### Spatial permutation testing

To assess the spatial alignment to previously characterized cortical maps, we employed the Spearman correlation coefficient (*rho*), which tested the association of between-hemisphere functional distance maps with the following public or homemade atlases: 1) the overall functional lateralization index across cognitive domains (*2*), 2) the evolutionary and developmental cortical expansion maps estimated by Hill and colleagues (*46*), and 3) the myelin (T1w/T2w) maps (*45*). The significance of the alignment was determined using a spatial spin test with 10,000 permutations, establishing a two-sided significance level (denoted as *p*_spin_) for assessing the statistical significance of *rho*. During the spin test, the brain maps were randomly rotated to maintain the spatial covariance structure of the original data, and the medial wall area was removed. This procedure was implemented by using the ‘spin_permutations’ function in BrainSpace. The Bonferroni method was applied to correct for the 4 comparisons, as *p*_spin_ < 0.05/4 = 0.0125 was considered to indicate statistical significance.

### Statistical analysis

Prior to the statistical analyses, 9 participants were excluded due to a lack of cognitive scores. We further identified 13 participants as outliers, whose values for at least one of the relevant variables (including global between-hemisphere functional distance, total brain volume, and cognitive scores) exceeded three standard deviations (SDs) from the corresponding mean of the entire group. Ultimately, a total of 755 participants were included in subsequent statistical analyses focused on individual differences. First, we tested the association between individual global between-hemisphere functional distance and demographics. In light of evidence that head motion is a substantial confound in functional connectivity analyses, although a series of head motion control and denoising techniques were applied during preprocessing, we also utilized the mean framewise displacement value across all runs as a covariate in our analysis to account for the potential effects of head movement. Partial correlation was employed to examine the relationships with age, handedness and total brain volume, while the F test was applied to compare the two sex groups (Fig. S6). To assess the association between global between-hemisphere functional distance and fluid composite score, we used partial correlations with age, sex, handedness, mean framewise displacement and total brain volume as baseline covariates. As a post hoc analysis, we conducted additional tests to explore the relationship between global between-hemisphere functional distance and each of the subdomain scores using partial correlation. Their significance was corrected using the Bonferroni method, with a significance threshold of *p* < 0.01 (0.05 divided by 5) deemed to indicate statistical significance.

To assess the stability of the association between global between-hemisphere functional distance and fluid intelligence, we estimated the sampling variability by calculating the distribution of the correlations in different subsamples (*47*). Specifically, we randomly selected participants with replacement from the full sample (n = 755) at equally spaced sample sizes (15 intervals; from 50 to 750). For each sample size, we randomly subsampled the participants 1,000 times. The sampling variability (95% confidence interval) at each sampling interval for both correlation coefficients and their significances are presented in Fig. S8.

The vertexwise between-hemisphere functional distances were initially smoothed with an 8mm FWHM kernel prior to statistical analysis; this was implemented by a workbench command (‘-metric-smoothing’) based on the group-averaged left mid-thickness surface. A general linear model (GLM) was applied on each vertex with age, sex, handedness, total brain volume and mean framewise displacement as covariates. Since we were interested only in brain regions that had a significant positive contribution to cognitive scores, we employed a one-tailed test. To correct the multiple vertexwise comparisons, the cortical vertices with false discovery rate (FDR)-corrected *p*_FDR_ < 0.05 were considered as significant. The statistical analysis of cortical vertices was performed by using the BrainStat toolbox (*83*).

To estimate the generalization of vertexwise prediction of fluid intelligence, leave-one-subject-out cross-validation (LOOCV) was used. In each iterative analysis, a linear support vector regression (SVR, C = 1) algorithm was trained as prediction mode using CANlab core functions (*84*) based on n − 1 participants, and the model is then tested on the remaining participant. Each participant was left out once. The performance of LOOCV was evaluated by calculating the Pearson coefficient between observed and predicted scores. To determine regions made reproducible contribution to the prediction model, bootstrap procedures with 5,000 samples (with replacement) were conducted.

### Mediation analysis

To determine whether between-hemisphere functional distance could play a mediating role in the effect of brain size on fluid intelligence/cognitive flexibility, we conducted a mediation analysis using the multilevel mediation and moderation (M3) toolbox (https://github.com/canlab/MediationToolbox). A mediation analysis tests whether the observed covariance between a predictor (X, brain size) and an outcome (Y, cognition) could be explained by a mediator (M, between-hemisphere functional distance). A significant mediation effect is obtained when the inclusion of M in a path model of the effect of X on Y significantly alters the slope of the X–Y relationship. Age, sex, handedness, and mean framewise displacement were included as covariates in the mediation model. This mediation model consists of four paths: (1) path c, the group effect on motor performance, that is, the total effect of the predictor on the outcome; (2) path a, the group effect on brain measures; (3) path b, the correlation between brain measures and motor scores, after controlling for the group factor; and (4) the a × b effect, which is referred to as the indirect effect and is indicative of whether the predictor-outcome relationship was significantly reduced after controlling for the mediator. We used bootstrapping (10,000 replacements) for significance testing (*85*).

### Validation analysis

To validate our main findings, we performed the analyses again by using the processed data after global signals regression, including both global and vertexwise correlation analyses and mediation analyses. We also conducted assessments of alternative hemispheric functional difference measures using different methodologies to substantiate the efficacy of our proposed framework. Specifically, we computed the homotopic FC as well as the lateralization of within-hemisphere FC strength. The homotopic FC was calculated by Pearson’s correlation between the time series of homotopic vertices, representing the temporal synchrony between hemispheres (*48*). The sum of positive values in the within-hemisphere FC profile was computed as within-hemispheric integration for each vertex. The lateralization of within-hemisphere FC strength was computed using the LI formula on within-hemispheric integration for each pair of homotopic vertices (*18*). As our focus was solely on the magnitude of hemispheric differences, we exclusively employed absolute values. For these two measures, we also examined their test-retest reliability and associations with behavioral fluid intelligence scores.

## Code and data availability

The code used in this paper is publicly available at https://github.com/liang-xinyu/Between-hemisphere-Functional-Divergence. The HCP rs-fMRI, and myelin data are publicly available (https://www.humanconnectome.org/). The calculated vertexwise hemispheric gradients are available at https://osf.io/pvdjq/.

## Acknowledgments

Thanks to Dr. Yunman Xia for discussing the framework and result interpretation.

## Author contributions

Conceptualization: XL, DV, EJ, GG Methodology: XL

Formal analysis: XL Investigation: XL, JL, LY Visualization: XL Supervision: GG Writing—original draft: XL

Writing—review & editing: XL, JL, LY, DV, EJ, GG

## Data and materials availability

All data needed to evaluate the conclusions in the paper are present in the paper and/or the Supplementary Materials. All data needed to evaluate the conclusions in the paper are present in the paper and the Supplementary Materials. The calculated vertexwise hemispheric gradients are available at https://osf.io/pvdjq/, and other generated cortical maps are available at https://github.com/liang-xinyu/Between-hemisphere-Functional-Divergence.

## References

1. P. Y. Herve, L. Zago, L. Petit, B. Mazoyer, N. Tzourio-Mazoyer, Revisiting human hemispheric specialization with neuroimaging. Trends Cogn Sci 17, 69–80 (2013).

2. V. R. Karolis, M. Corbetta, M. Thiebaut de Schotten, The architecture of functional lateralisation and its relationship to callosal connectivity in the human brain. Nat Commun 10, 1417 (2019).

3. R. Sperry, Some effects of disconnecting the cerebral hemispheres. Science 217, 1223–6 (1982).

4. A. W. Toga, P. M. Thompson, Mapping brain asymmetry. Nat Rev Neurosci 4, 37–48 (2003).

5. S. J. Gotts, H. J. Jo, G. L. Wallace, Z. S. Saad, R. W. Cox, A. Martin, Two distinct forms of functional lateralization in the human brain. Proc Natl Acad Sci U S A 110, E3435–44 (2013).

6. D. Wang, R. L. Buckner, H. Liu, Functional specialization in the human brain estimated by intrinsic hemispheric interaction. J Neurosci 34, 12341–52 (2014).

7. O. Güntürkün, S. Ocklenburg, Ontogenesis of Lateralization. Neuron 94, 249–263 (2017).

8. K. Hugdahl, Hemispheric asymmetry: contributions from brain imaging. WIREs Cognitive Science 2, 461–478 (2011).

9. J. L. Ringo, R. W. Doty, S. Demeter, P. Y. Simard, Time is of the essence: a conjecture that hemispheric specialization arises from interhemispheric conduction delay. Cereb Cortex 4, 331–43 (1994).

10. G. Vallortigara, L. J. Rogers, Survival with an asymmetrical brain: Advantages and disadvantages of cerebral lateralization. Behavioral and Brain Sciences 28, 575–589 (2005).

11. D. V. Bishop, Cerebral asymmetry and language development: cause, correlate, or consequence? Science 340, 1230531 (2013).

12. O. Güntürkün, F. Ströckens, S. Ocklenburg, Brain Lateralization: A Comparative Perspective. Physiol Rev 100, 1019–1063 (2020).

13. G. Hartwigsen, Y. Bengio, D. Bzdok, How does hemispheric specialization contribute to human-defining cognition? Neuron, S0896627321002907 (2021).

14. R. B. Cattell, Theory of fluid and crystallized intelligence: A critical experiment. Journal of Educational Psychology 54, 1–22 (1963).

15. J. Levy, The Mammalian Brain and the Adaptive Advantage of Cerebral Asymmetry. Annals of the New York Academy of Sciences 299, 264–272 (1977).

16. S. Dimond, G. Beaumont, Use of Two Cerebral Hemispheres to increase Brain Capacity. Nature 232, 270–271 (1971).

17. G. Vallortigara, L. J. Rogers, A function for the bicameral mind. Cortex 124, 274–285 (2020).

18. M. Joliot, N. Tzourio-Mazoyer, B. Mazoyer, Intra-hemispheric intrinsic connectivity asymmetry and its relationships with handedness and language Lateralization. Neuropsychologia 93, 437–447 (2016).

19. H. Liu, S. M. Stufflebeam, J. Sepulcre, T. Hedden, R. L. Buckner, Evidence from intrinsic activity that asymmetry of the human brain is controlled by multiple factors. P Natl Acad Sci USA 106, 20499–503 (2009).

20. H. J. Jo, Z. S. Saad, S. J. Gotts, A. Martin, R. W. Cox, Quantifying agreement between anatomical and functional interhemispheric correspondences in the resting brain. PLoS One 7, e48847 (2012).

21. J. V. Haxby, J. S. Guntupalli, S. A. Nastase, M. Feilong, Hyperalignment: Modeling shared information encoded in idiosyncratic cortical topographies. eLife 9, e56601 (2020).

22. J. S. Guntupalli, M. Feilong, J. V. Haxby, A computational model of shared fine-scale structure in the human connectome. PLOS Computational Biology 14, e1006120 (2018).

23. X. Liang, C. Zhao, X. Jin, Y. Jiang, L. Yang, Y. Chen, G. Gong, Sex-related human brain asymmetry in hemispheric functional gradients. NeuroImage, 117761 (2021).

24. T. R. del J. Gonzalez Alam, B. L. A. Mckeown, Z. Gao, B. Bernhardt, R. Vos de Wael, D. S. Margulies, J. Smallwood, E. Jefferies, A tale of two gradients: differences between the left and right hemispheres predict semantic cognition. Brain Struct Funct, doi: 10.1007/s00429-021-02374-w (2021).

25. B. Wan, Ş. Bayrak, T. Xu, H. L. Schaare, R. A. I. Bethlehem, B. C. Bernhardt, S. L. Valk, “Asymmetry of cortical functional hierarchy in humans and macaques suggests phylogenetic conservation and adaptation” (2021); 10.1101/2021.11.03.466058.

26. D. S. Margulies, S. Ovadia-Caro, N. Saadon-Grosman, B. Bernhardt, B. Jefferies, J. Smallwood, “Cortical Gradients and Their Role in Cognition” in Encyclopedia of Behavioral Neuroscience, 2nd Edition (Elsevier, 2022; https://linkinghub.elsevier.com/retrieve/pii/B9780128196410000104), pp. 242–250.

27. J. M. Huntenburg, P. L. Bazin, D. S. Margulies, Large-Scale Gradients in Human Cortical Organization. Trends Cogn Sci 22, 21–31 (2018).

28. D. C. Van Essen, S. M. Smith, D. M. Barch, T. E. Behrens, E. Yacoub, K. Ugurbil, W. U-Minn HCP Consortium, The WU-Minn Human Connectome Project: an overview. Neuroimage 80, 62–79 (2013).

29. S. R. Cox, S. J. Ritchie, C. Fawns-Ritchie, E. M. Tucker-Drob, I. J. Deary, Structural brain imaging correlates of general intelligence in UK Biobank. Intelligence 76, 101376 (2019).

30. J. Pietschnig, L. Penke, J. M. Wicherts, M. Zeiler, M. Voracek, Meta-analysis of associations between human brain volume and intelligence differences: How strong are they and what do they mean? Neuroscience & Biobehavioral Reviews 57, 411–432 (2015).

31. D. van der Linden, C. S. Dunkel, G. Madison, Sex differences in brain size and general intelligence (g). Intelligence 63, 78–88 (2017).

32. X. Kang, T. J. Herron, M. Ettlinger, D. L. Woods, Hemispheric asymmetries in cortical and subcortical anatomy. Laterality 20, 658–684 (2015).

33. X.-Z. Kong, S. R. Mathias, T. Guadalupe, E. L. W. Group, D. C. Glahn, B. Franke, F. Crivello, N. Tzourio-Mazoyer, S. E. Fisher, P. M. Thompson, C. Francks, Mapping cortical brain asymmetry in 17,141 healthy individuals worldwide via the ENIGMA Consortium. PNAS 115, E5154–E5163 (2018).

34. N. Tzourio-Mazoyer, L. Petit, A. Razafimandimby, F. Crivello, L. Zago, G. Jobard, M. Joliot, E. Mellet, B. Mazoyer, Left Hemisphere Lateralization for Language in Right-Handers Is Controlled in Part by Familial Sinistrality, Manual Preference Strength, and Head Size. J. Neurosci. 30, 13314–13318 (2010).

35. S. F. Witelson, H. Beresh, D. L. Kigar, Intelligence and brain size in 100 postmortem brains: sex, lateralization and age factors. Brain 129, 386–398 (2006).

36. L. J. Rogers, G. Vallortigara, When and Why Did Brains Break Symmetry? Symmetry 7, 2181–2194 (2015).

37. R. Vos de Wael, O. Benkarim, C. Paquola, S. Lariviere, J. Royer, S. Tavakol, T. Xu, S. J. Hong, G. Langs, S. Valk, B. Misic, M. Milham, D. Margulies, J. Smallwood, B. C. Bernhardt, BrainSpace: a toolbox for the analysis of macroscale gradients in neuroimaging and connectomics datasets. Commun Biol 3, 103 (2020).

38. B. T. Yeo, F. M. Krienen, J. Sepulcre, M. R. Sabuncu, D. Lashkari, M. Hollinshead, J. L. Roffman, J. W. Smoller, L. Zollei, J. R. Polimeni, B. Fischl, H. Liu, R. L. Buckner, The organization of the human cerebral cortex estimated by intrinsic functional connectivity. J Neurophysiol 106, 1125–65 (2011).

39. L. Yang, C. Zhao, Y. Xiong, S. Zhong, D. Wu, S. Peng, M. T. de Schotten, G. Gong, Callosal Fiber Length Scales with Brain Size According to Functional Lateralization, Evolution, and Development. J. Neurosci. 42, 3599–3610 (2022).

40. D. S. Margulies, S. S. Ghosh, A. Goulas, M. Falkiewicz, J. M. Huntenburg, G. Langs, G. Bezgin, S. B. Eickhoff, F. X. Castellanos, M. Petrides, E. Jefferies, J. Smallwood, Situating the default-mode network along a principal gradient of macroscale cortical organization. Proc Natl Acad Sci USA 113, 12574–12579 (2016).

41. T. Yarkoni, R. A. Poldrack, T. E. Nichols, D. C. Van Essen, T. D. Wager, Large-scale automated synthesis of human functional neuroimaging data. Nat Methods 8, 665–670 (2011).

42. J. S. Siegel, A. Mitra, T. O. Laumann, B. A. Seitzman, M. Raichle, M. Corbetta, A. Z. Snyder, Data Quality Influences Observed Links Between Functional Connectivity and Behavior. Cerebral Cortex 27, 4492–4502 (2017).

43. N. Akshoomoff, J. L. Beaumont, P. J. Bauer, S. S. Dikmen, R. C. Gershon, D. Mungas, J. Slotkin, D. Tulsky, S. Weintraub, P. D. Zelazo, R. K. Heaton, Viii. Nih Toolbox Cognition Battery (cb): Composite Scores of Crystallized, Fluid, and Overall Cognition. Monographs of the Society for Research in Child Development 78, 119–132 (2013).

44. M. F. Glasser, T. S. Coalson, E. C. Robinson, C. D. Hacker, J. Harwell, E. Yacoub, K. Ugurbil, J. Andersson, C. F. Beckmann, M. Jenkinson, S. M. Smith, D. C. Van Essen, A multi-modal parcellation of human cerebral cortex. Nature 536, 171–178 (2016).

45. M. F. Glasser, D. C. V. Essen, Mapping Human Cortical Areas In Vivo Based on Myelin Content as Revealed by T1-and T2-Weighted MRI. J. Neurosci. 31, 11597–11616 (2011).

46. J. Hill, T. Inder, J. Neil, D. Dierker, J. Harwell, D. V. Essen, Similar patterns of cortical expansion during human development and evolution. PNAS 107, 13135–13140 (2010).

47. S. Marek, B. Tervo-Clemmens, F. J. Calabro, D. F. Montez, B. P. Kay, A. S. Hatoum, M. R. Donohue, W. Foran, R. L. Miller, T. J. Hendrickson, S. M. Malone, S. Kandala, E. Feczko, O. Miranda-Dominguez, A. M. Graham, E. A. Earl, A. J. Perrone, M. Cordova, O. Doyle, L. A. Moore, G. M. Conan, J. Uriarte, K. Snider, B. J. Lynch, J. C. Wilgenbusch, T. Pengo, A. Tam, J. Chen, D. J. Newbold, A. Zheng, N. A. Seider, A. N. Van, A. Metoki, R. J. Chauvin, T. O. Laumann, D. J. Greene, S. E. Petersen, H. Garavan, W. K. Thompson, T. E. Nichols, B. T. T. Yeo, D. M. Barch, B. Luna, D. A. Fair, N. U. F. Dosenbach, Reproducible brain-wide association studies require thousands of individuals. Nature 603, 654–660 (2022).

48. X. Jin, X. Liang, G. Gong, Functional Integration Between the Two Brain Hemispheres: Evidence From the Homotopic Functional Connectivity Under Resting State. Front. Neurosci. 14, 932 (2020).

49. Z. Gracia-Tabuenca, M. B. Moreno, F. A. Barrios, S. Alcauter, Hemispheric asymmetry and homotopy of resting state functional connectivity correlate with visuospatial abilities in school-age children. Neuroimage 174, 441–448 (2018).

50. Y. Sun, J. Li, J. Suckling, L. Feng, Asymmetry of Hemispheric Network Topology Reveals Dissociable Processes between Functional and Structural Brain Connectome in Community-Living Elders. Front Aging Neurosci 9, 361 (2017).

51. L. Tian, J. Wang, C. Yan, Y. He, Hemisphere- and gender-related differences in small-world brain networks: a resting-state functional MRI study. Neuroimage 54, 191–202 (2011).

52. N. Tzourio-Mazoyer, “Intra- and Inter-hemispheric Connectivity Supporting Hemispheric Specialization” in Micro-, Meso- and Macro-Connectomics of the Brain, H. Kennedy, D. C. Van Essen, Y. Christen, Eds. (Cham (CH), 2016), pp. 129–146.

53. A. R. Bradshaw, D. V. M. Bishop, Z. V. J. Woodhead, Methodological considerations in assessment of language lateralisation with fMRI: a systematic review. PeerJ 5, e3557 (2017).

54. M. L. Seghier, Categorical laterality indices in fMRI: a parallel with classic similarity indices. Brain Struct Funct 224, 1377–1383 (2019).

55. J. R. Gray, C. F. Chabris, T. S. Braver, Neural mechanisms of general fluid intelligence. Nat Neurosci 6, 316–322 (2003).

56. M. L. Dixon, A. De La Vega, C. Mills, J. Andrews-Hanna, R. N. Spreng, M. W. Cole, K. Christoff, Heterogeneity within the frontoparietal control network and its relationship to the default and dorsal attention networks. Proceedings of the National Academy of Sciences 115, E1598–E1607 (2018).

57. X. Wang, K. Krieger-Redwood, B. Lyu, R. Lowndes, G. Wu, N. E. Souter, X. Wang, R. Kong, G. Shafiei, B. C. Bernhardt, Z. Cui, J. Smallwood, Y. Du, E. Jefferies, “The brain’s topographical organization shapes dynamic interaction patterns to support flexible behavior” (preprint, Neuroscience, 2023); 10.1101/2023.09.06.556465.

58. R. E. Jung, R. J. Haier, The Parieto-Frontal Integration Theory (P-FIT) of intelligence: Converging neuroimaging evidence. Behavioral and Brain Sciences 30, 135–154 (2007).

59. J. Duncan, The multiple-demand (MD) system of the primate brain: mental programs for intelligent behaviour. Trends in Cognitive Sciences 14, 172–179 (2010).

60. M. Assem, M. F. Glasser, D. C. Van Essen, J. Duncan, A Domain-General Cognitive Core Defined in Multimodally Parcellated Human Cortex. Cerebral Cortex 30, 4361–4380 (2020).

61. A. K. Barbey, Network Neuroscience Theory of Human Intelligence. Trends in Cognitive Sciences 22, 8–20 (2018).

62. G. Vallortigara, L. J., Rogers survival with an asymmetrical brain: advantages and disadvantages of cerebral lateralization. Behavioral and Brain Sciences 28, 575–589 (2005).

63. L. J. Rogers, Brain Lateralization and Cognitive Capacity. Animals 11, 1996 (2021).

64. F. Kurth, P. M. Thompson, E. Luders, Investigating the differential contributions of sex and brain size to gray matter asymmetry. Cortex 99, 235–242 (2018).

65. R. L. Buckner, F. M. Krienen, The evolution of distributed association networks in the human brain. Trends Cogn Sci 17, 648–65 (2013).

66. J. K. Rilling, Comparative primate neurobiology and the evolution of brain language systems. Current Opinion in Neurobiology 28, 10–14 (2014).

67. J. Hänggi, L. Fövenyi, F. Liem, M. Meyer, L. Jäncke, The hypothesis of neuronal interconnectivity as a function of brain size—a general organization principle of the human connectome. Frontiers in Human Neuroscience 8 (2014).

68. R. K. Heaton, N. Akshoomoff, D. Tulsky, D. Mungas, S. Weintraub, S. Dikmen, J. Beaumont, K. B. Casaletto, K. Conway, J. Slotkin, R. Gershon, Reliability and Validity of Composite Scores from the NIH Toolbox Cognition Battery in Adults. Journal of the International Neuropsychological Society 20, 588–598 (2014).

69. M. F. Glasser, S. N. Sotiropoulos, J. A. Wilson, T. S. Coalson, B. Fischl, J. L. Andersson, J. Xu, S. Jbabdi, M. Webster, J. R. Polimeni, D. C. Van Essen, M. Jenkinson, W. U-Minn HCP Consortium, The minimal preprocessing pipelines for the Human Connectome Project. Neuroimage 80, 105–24 (2013).

70. L. Griffanti, G. Salimi-Khorshidi, C. F. Beckmann, E. J. Auerbach, G. Douaud, C. E. Sexton, E. Zsoldos, K. P. Ebmeier, N. Filippini, C. E. Mackay, S. Moeller, J. Xu, E. Yacoub, G. Baselli, K. Ugurbil, K. L. Miller, S. M. Smith, ICA-based artefact removal and accelerated fMRI acquisition for improved resting state network imaging. Neuroimage 95, 232–47 (2014).

71. G. Salimi-Khorshidi, G. Douaud, C. F. Beckmann, M. F. Glasser, L. Griffanti, S. M. Smith, Automatic denoising of functional MRI data: combining independent component analysis and hierarchical fusion of classifiers. Neuroimage 90, 449–68 (2014).

72. S. M. Smith, C. F. Beckmann, J. Andersson, E. J. Auerbach, J. Bijsterbosch, G. Douaud, E. Duff, D. A. Feinberg, L. Griffanti, M. P. Harms, M. Kelly, T. Laumann, K. L. Miller, S. Moeller, S. Petersen, J. Power, G. Salimi-Khorshidi, A. Z. Snyder, A. T. Vu, M. W. Woolrich, J. Xu, E. Yacoub, K. Ugurbil, D. C. Van Essen, M. F. Glasser, W. U-Minn HCP Consortium, Resting-state fMRI in the Human Connectome Project. Neuroimage 80, 144–68 (2013).

73. M. J. Lowe, D. P. Russell, Treatment of baseline drifts in fMRI time series analysis. J Comput Assist Tomogr 23, 463–73 (1999).

74. K. Murphy, M. D. Fox, Towards a consensus regarding global signal regression for resting state functional connectivity MRI. Neuroimage 154, 169–173 (2017).

75. D. Cordes, V. M. Haughton, K. Arfanakis, J. D. Carew, P. A. Turski, C. H. Moritz, M. A. Quigley, M. E. Meyerand, Frequencies contributing to functional connectivity in the cerebral cortex in “resting-state” data. AJNR Am J Neuroradiol 22, 1326–33 (2001).

76. J. D. Power, A. Mitra, T. O. Laumann, A. Z. Snyder, B. L. Schlaggar, S. E. Petersen, Methods to detect, characterize, and remove motion artifact in resting state fMRI. NeuroImage 84, 320–341 (2014).

77. J. W. Cho, A. Korchmaros, J. T. Vogelstein, M. P. Milham, T. Xu, Impact of concatenating fMRI data on reliability for functional connectomics. NeuroImage 226, 117549 (2021).

78. S. J. Hong, R. Vos de Wael, R. A. I. Bethlehem, S. Lariviere, C. Paquola, S. L. Valk, M. P. Milham, A. Di Martino, D. S. Margulies, J. Smallwood, B. C. Bernhardt, Atypical functional connectome hierarchy in autism. Nat Commun 10, 1022 (2019).

79. R. R. Coifman, S. Lafon, A. B. Lee, M. Maggioni, B. Nadler, F. Warner, S. W. Zucker, Geometric diffusions as a tool for harmonic analysis and structure definition of data: diffusion maps. Proc Natl Acad Sci U S A 102, 7426–31 (2005).

80. A. H. Williams, E. Kunz, S. Kornblith, S. W. Linderman, Generalized Shape Metrics on Neural Representations. arXiv arXiv:2110.14739 [Preprint] (2022). http://arxiv.org/abs/2110.14739.

81. R. A. I. Bethlehem, C. Paquola, J. Seidlitz, L. Ronan, B. Bernhardt, C. C. Consortium, K. A. Tsvetanov, Dispersion of functional gradients across the adult lifespan. Neuroimage, 117299 (2020).

82. R. A. Poldrack, J. A. Mumford, T. Schonberg, D. Kalar, B. Barman, T. Yarkoni, Discovering Relations Between Mind, Brain, and Mental Disorders Using Topic Mapping. PLOS Computational Biology 8, e1002707 (2012).

83. K. Worsley, J. Taylor, F. Carbonell, M. Chung, E. Duerden, B. Bernhardt, O. Lyttelton, M. Boucher, A. Evans, SurfStat: A Matlab toolbox for the statistical analysis of univariate and multivariate surface and volumetric data using linear mixed effects models and random field theory. NeuroImage 47, S102 (2009).

84. L. Kohoutová, J. Heo, S. Cha, S. Lee, T. Moon, T. D. Wager, C.-W. Woo, Toward a unified framework for interpreting machine-learning models in neuroimaging. Nat Protoc 15, 1399–1435 (2020).

85. P. E. Shrout, N. Bolger, Mediation in experimental and nonexperimental studies: New procedures and recommendations. Psychological Methods 7, 422–445 (2002).

