## Supplementary Materials for "Functional divergence between the two cerebral hemispheres contributes to human fluid intelligence"

Xinyu Liang^*^ *et al.*

**This PDF file includes:**

Supplementary Text

Figs. S1 to S11

Tables S1 to S4

Supplementary Text

Contribution of each gradient to between-hemisphere functional distance

To determine which gradient contributes the most to between-hemisphere functional distance, we estimated the correlation between the hemispheric difference within each single gradient and the proposed functional distance (Fig. S3). For the similarity of spatial distribution, the hemispheric difference in the principal gradient (*r* = 0.77 ± 0.06) made the greatest contribution to the between-hemisphere functional distance, while the hemispheric difference in the tertiary gradient was also highly correlated with the between-hemisphere functional distance (*r* = 0.71 ± 0.06). Regarding to the global measurements, both the overall hemispheric differences of the tertiary (*r* = 0.68, *p* < 0.001) and the principal gradients (*r* = 0.65, *p* < 0.001) showed very high correlations with the global between-hemisphere functional distance. In addition, only the secondary gradient had a relatively low contribution (spatial correlation: *r* = 0.37 ± 0.10; global correlation: *r* = 0.32, *p* < 0.001) to the between-hemisphere functional distance among all 6 gradients.

Validation results with global signal regression

Given the debates regarding the influences of GS, we performed additional validation analyses based on the processed data after GS regression. The results showed a very consistent spatial pattern of group-averaged cortical between-hemisphere functional distance and its test-retest reliability. We also found that the individual global between-hemisphere functional distance was related to fluid intelligences (Table S1) and brain size (Table S2). We successfully built a significant mediation pathway in which brain size, global between-hemisphere functional distance, and fluid cognition composites acted as the predictor, mediator, and outcome, respectively (Fig. S8d). To identify brain regions that were significantly associated with fluid composite scores, we performed the same GLM analysis on cortical vertices. The resultant statistical map exhibited a very similar pattern to our main findings (Fig. S8e). We found fewer significant regions, while the same associations between fluid intelligence and between-hemisphere functional distance were found in the inferior parietal cortex (area PFm), anterior dorsolateral frontal cortex (area a9-46v), anterior ventral insular (area AVI), and dorsal part of the ventral anterior cingulate cortex (area 24dd).

Comparison with other functional lateralization measures

The group-level spatial patterns of homotopic FC and lateralization of within-hemisphere FC strength are shown in Fig. S7. Their spatial patterns showed that the hemispheric differences varied across cortical regions, indicating a similar distribution to our proposed between-hemisphere functional distance. We also evaluated the test-retest reliability between the two sessions. The results showed that homotopic FC displayed an ICC comparable to that of between-hemisphere functional distance (global = 0.68, vertexwise = 0.60 ± 0.14, Fig. S11a), whereas the ICC of lateralization of within-hemisphere FC strength was much lower (global = 0.52, vertexwise = 0.18 ± 0.07, Fig. S11b). Subsequently, we replicated the correlation analyses between these two forms of hemispheric functional differences and cognitive abilities. This analysis was carried out to determine whether these alternative measures also exhibited substantial effects on fluid intelligence. Notably, neither global metric demonstrated statistically significant correlations with the fluid cognitive composite or its subdomain scores (Table S3). Collectively, our results suggested that between-hemisphere functional distance is not only a more reliable measure but also reflects more individual variance related to actual cognitive outcomes.

Details of cortical regions related to fluid intelligence

To further identify the attribution of the significant vertices related to fluid intelligence, we located them by using Yeo’s symmetric networks and multimodal parcellation. At the network level, one-third of the significant vertices are situated in the divergent zone, and another one-third of the vertices belonged to the FPN (Fig. S7). This highlighted the significant role of the divergent zone and control network in fluid cognitive processing. Due to the asymmetric distribution of regions in multimodal parcellation, then we projected the significant vertices to the left and right hemispheres. The results showed that these vertices were mainly located in the PFm and the a9-46v areas for both hemispheres (Table S4).

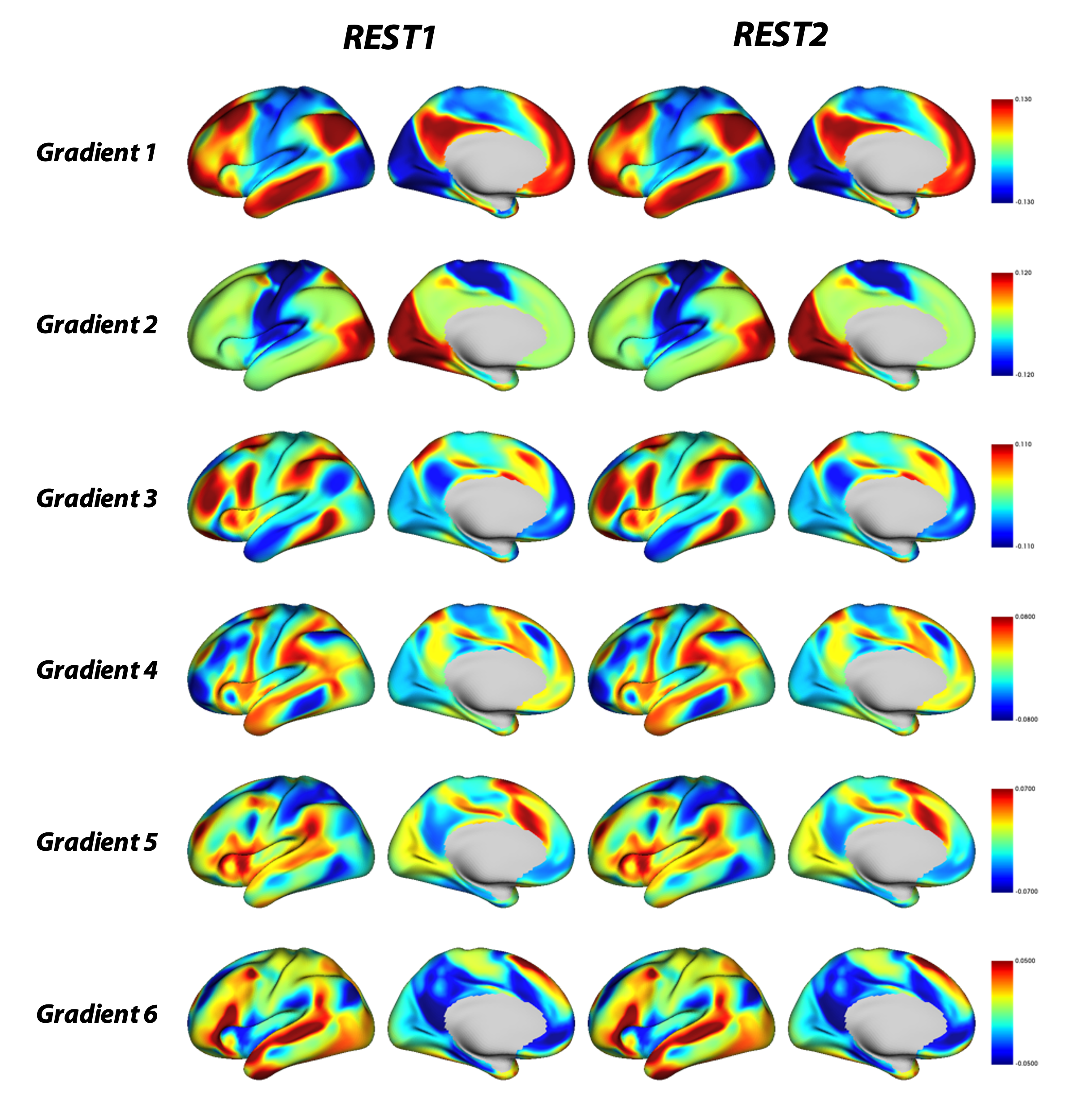

Fig. S1. Hemispheric gradient templates utilized for hyperalignment.

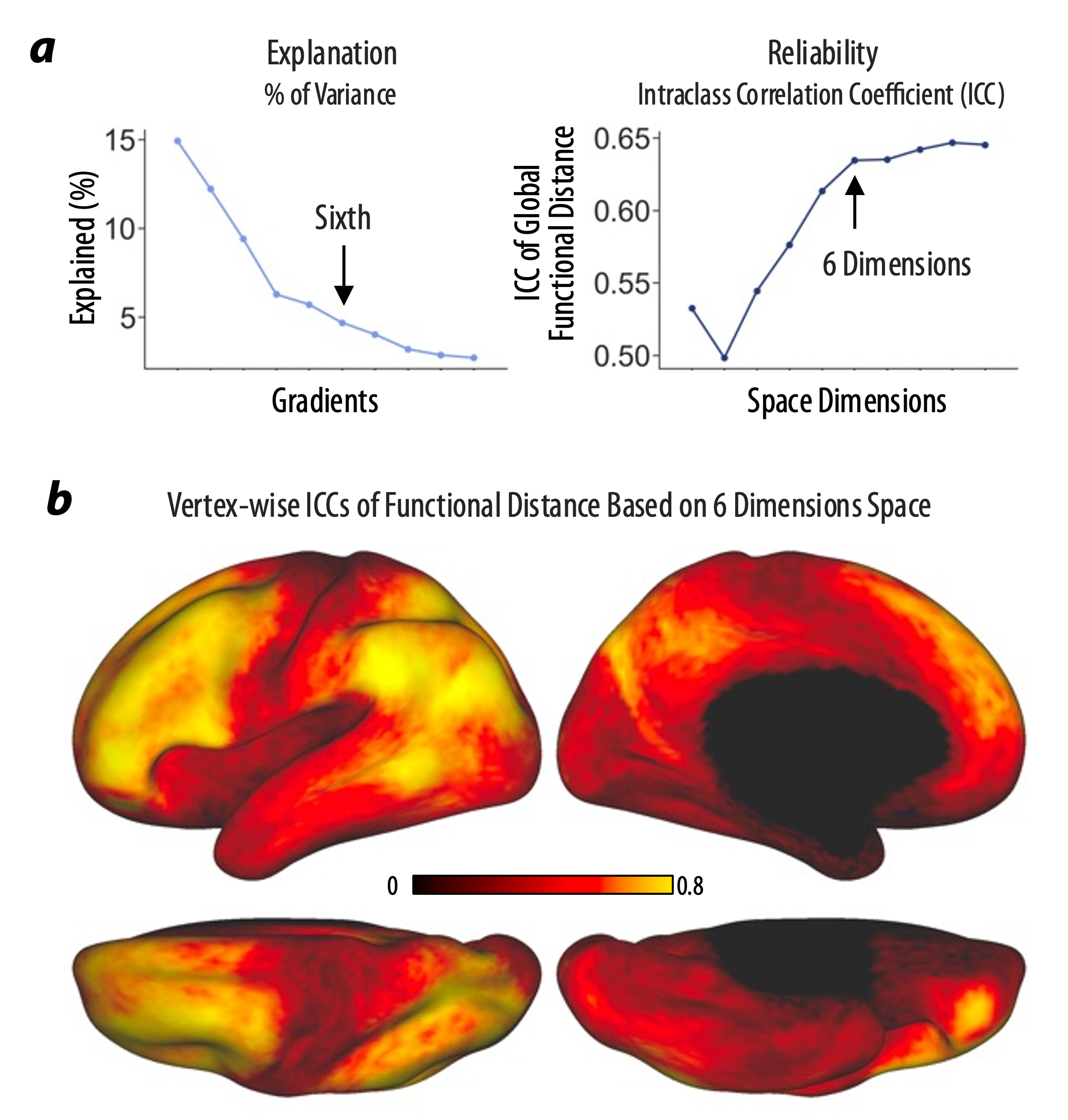

Fig. S2. Dimension selection and vertexwise reliability of between-hemisphere functional distance based on 6-dimensional space.

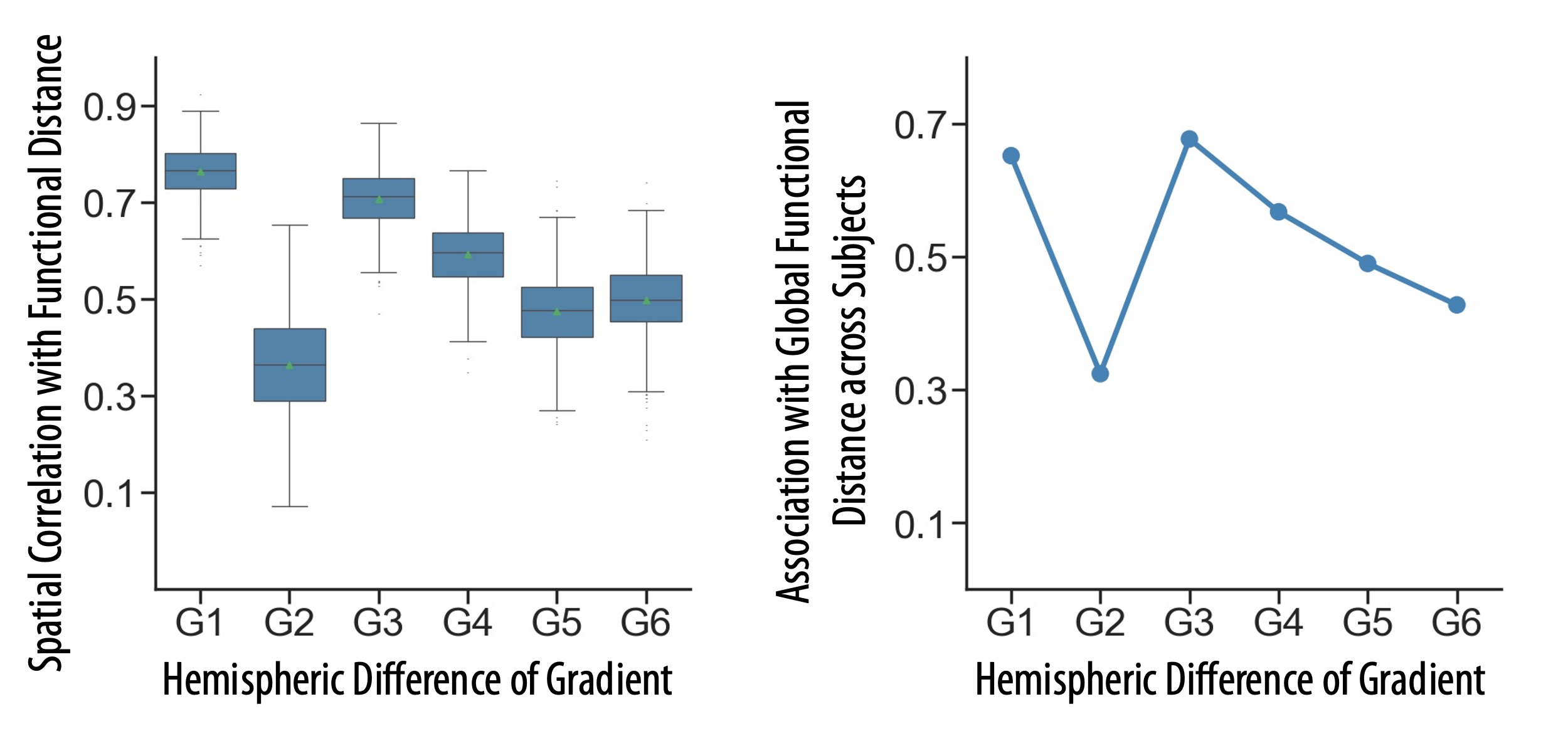

Fig. S3. Contributions of each gradient to between-hemisphere functional distance. To determine which gradient has a high contribution to between-hemisphere functional distance, we estimated the correlation between the hemispheric difference within each single gradient and the proposed functional distance. The left plot shows the individual distribution of spatial similarity, and the right plot shows the similarity of global hemispheric distances across participants.

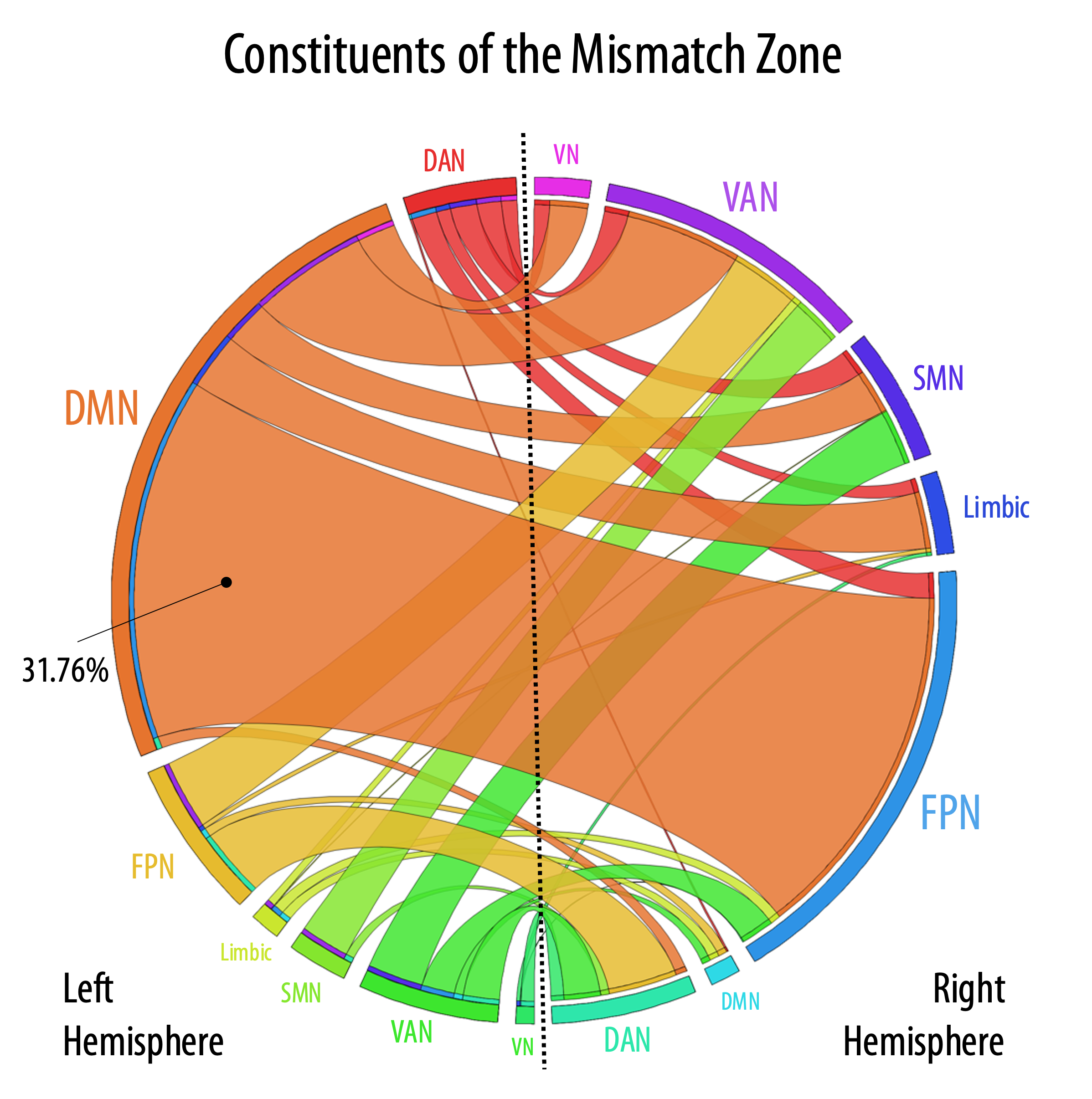

Fig. S4. The constituents of the mismatch zone in Yeo’s 7 networks. Owing to the asymmetric distribution of Yeo’s 7 networks, we designated a mismatch zone in which homotopic vertices belong to different networks across the two hemispheres. The vertices belonging to the DMN in the left hemisphere and the vertices belonging to the FPN in the right hemisphere contributed the most to this mismatch zone (accounting for 31.76%). Abbreviations: FPN, frontoparietal control network; DAN, dorsal attention network; DMN, default mode network; VAN, ventral attention network; SMN, somatomotor network; and VN, visual network.

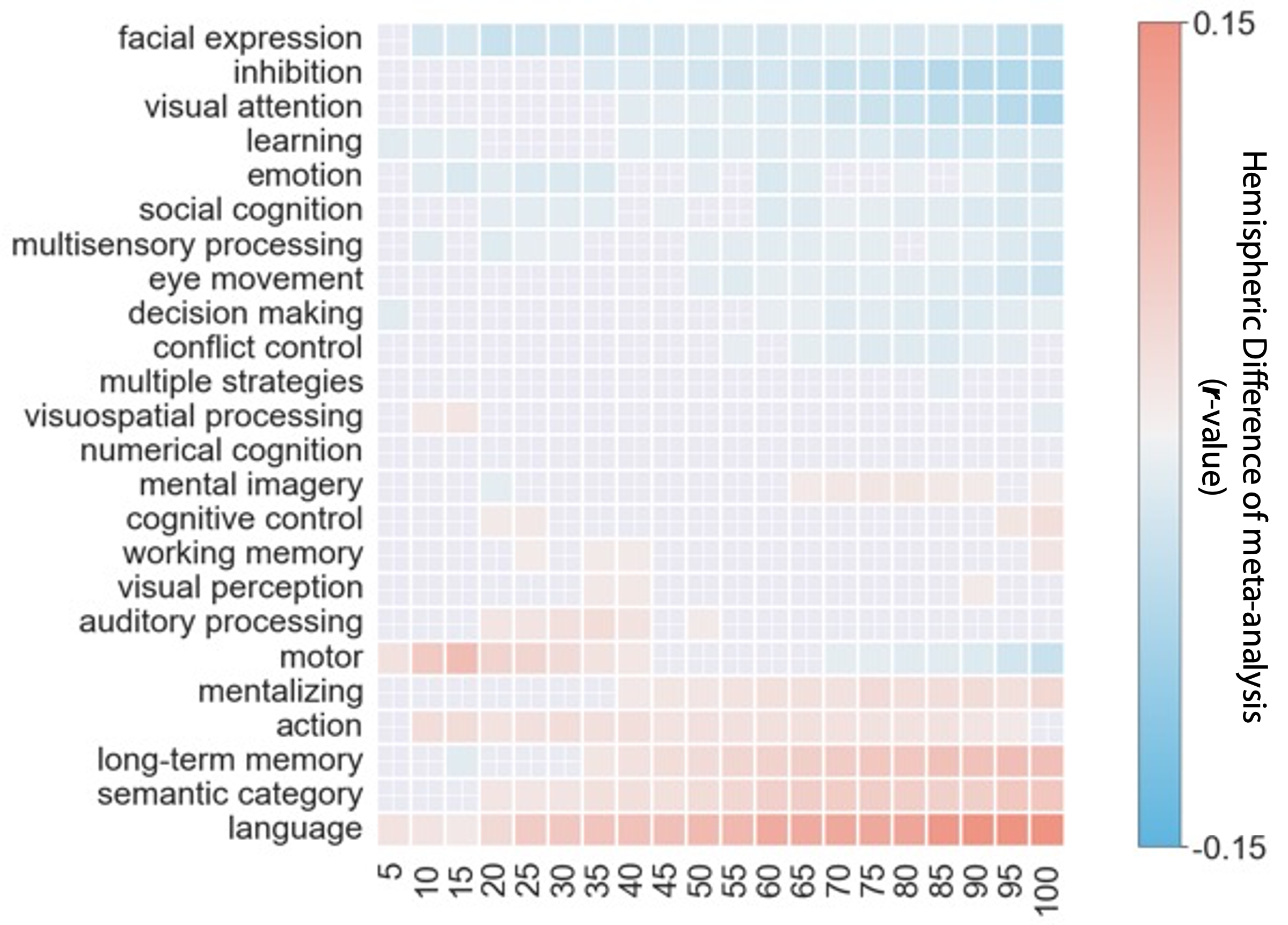

Fig. S5. Hemispheric difference in the meta-analysis results. To visualize the hemispheric difference between topic term-based decoding results from the left and right hemispheres, we calculated the deviation of the correction coefficient (left–right) for each topic term and each 5- percentile bin. The order of the terms follows an ascending trend according to the average of differences across all bins, suggesting a general transition from rightward to leftward lateralization. The arbitrary threshold is set as 0.01 for visualization.

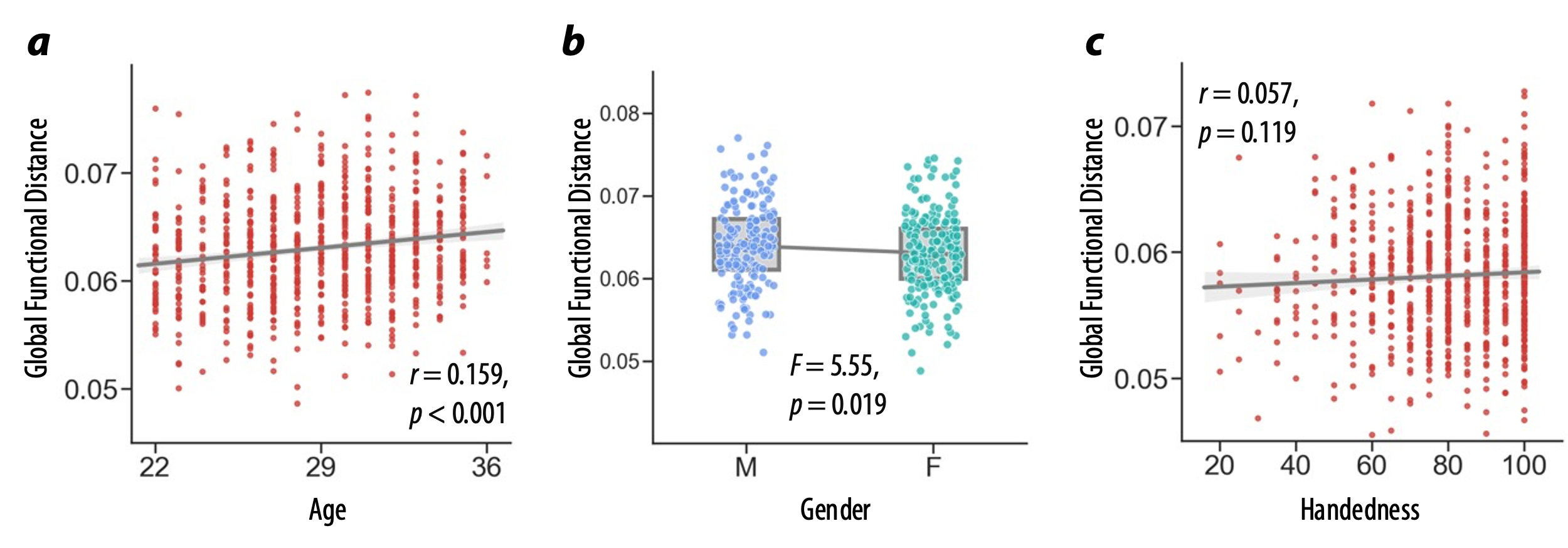

Fig. S6. The associations between individual global between-hemisphere functional distance and demographics.

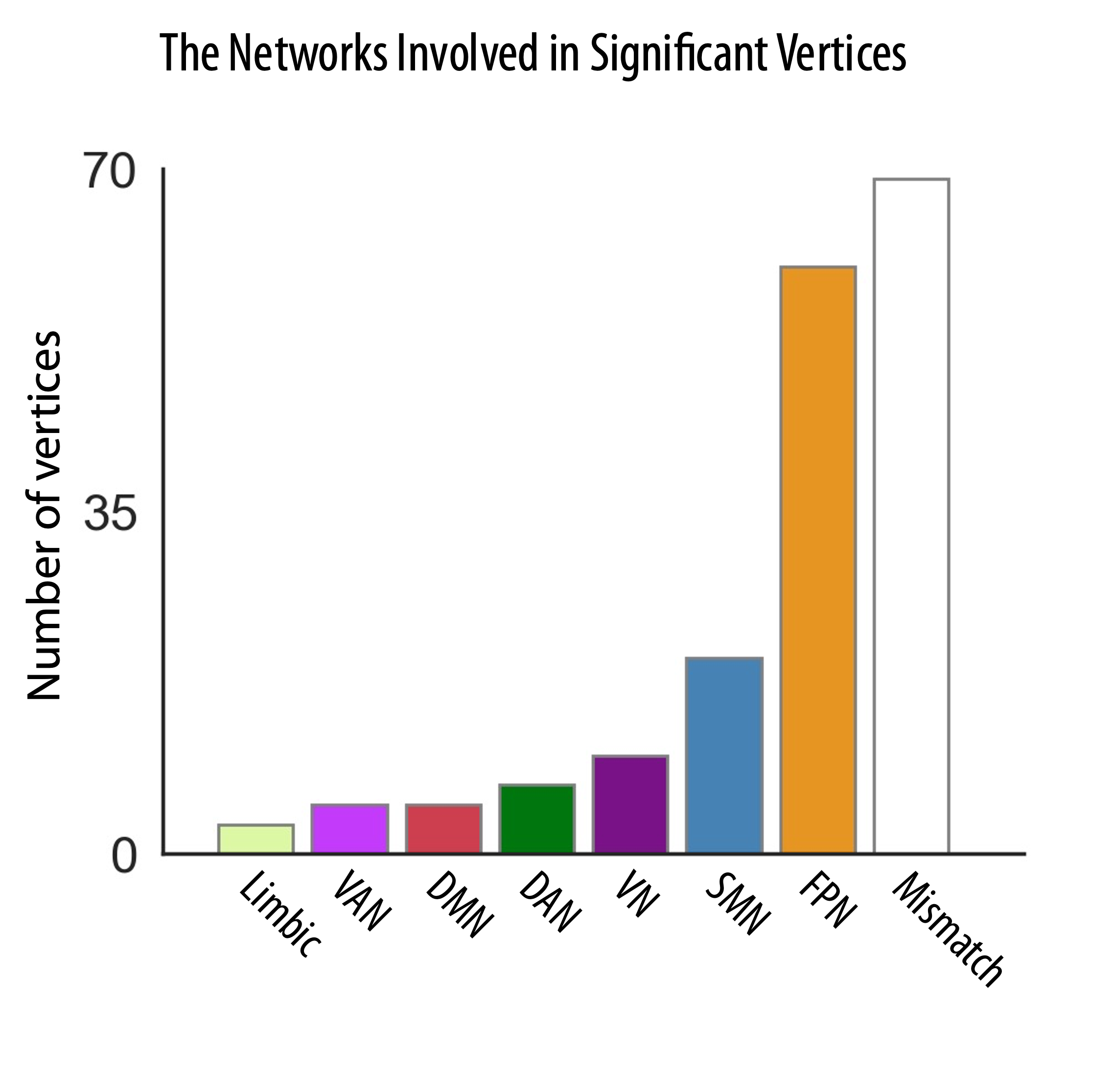

Fig. S7. The networks involved in the association between vertexwise between-hemisphere functional distance and fluid intelligence.

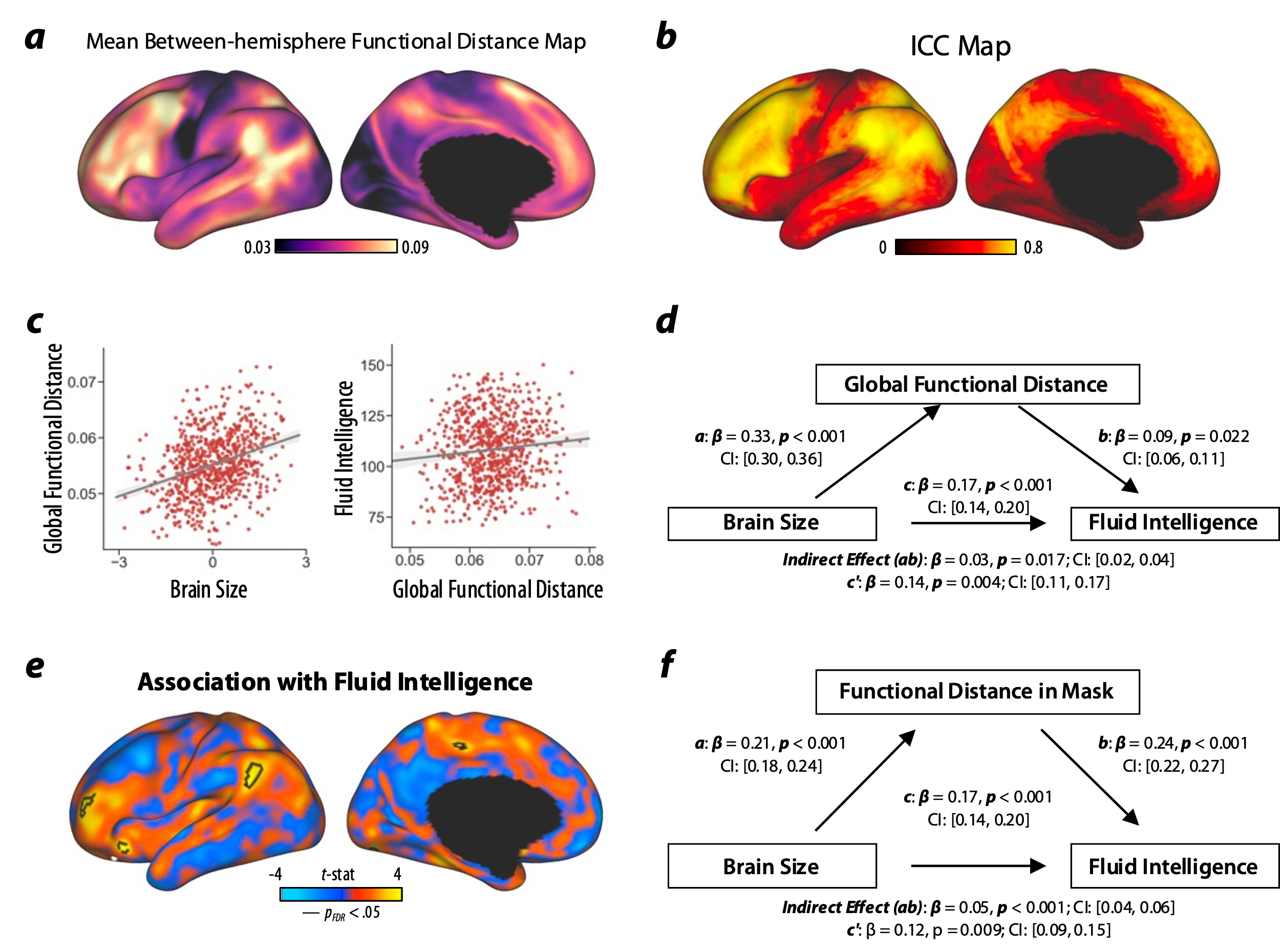

Fig. S8. The validation results based on preprocessed data after global signal regression.

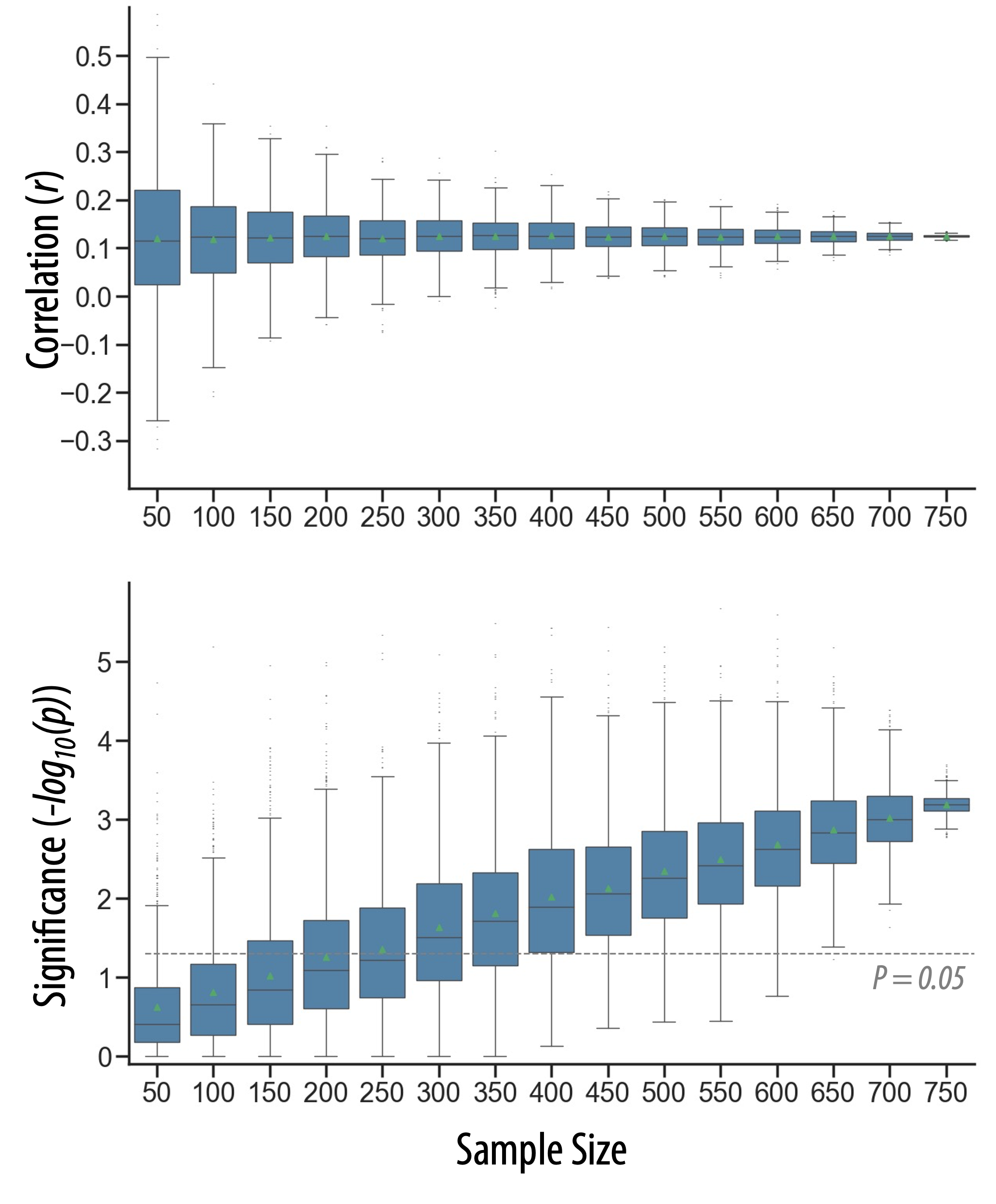

Fig. S9. The sampling variability of associations between global between-hemisphere functional distance and fluid intelligence. We estimated the influence of sample size on the stability of the association. We computed sampling variability for a range of sampling rates (15 intervals evenly spaced from 50 to 750). In each interval, we randomly selected participants with replacement from the full sample (n = 755) to obtain 1000 subsamples. We plot the distribution of similarity and significance across all subsamples. The results show that the association becomes significant and stable when the sample size reaches 650.

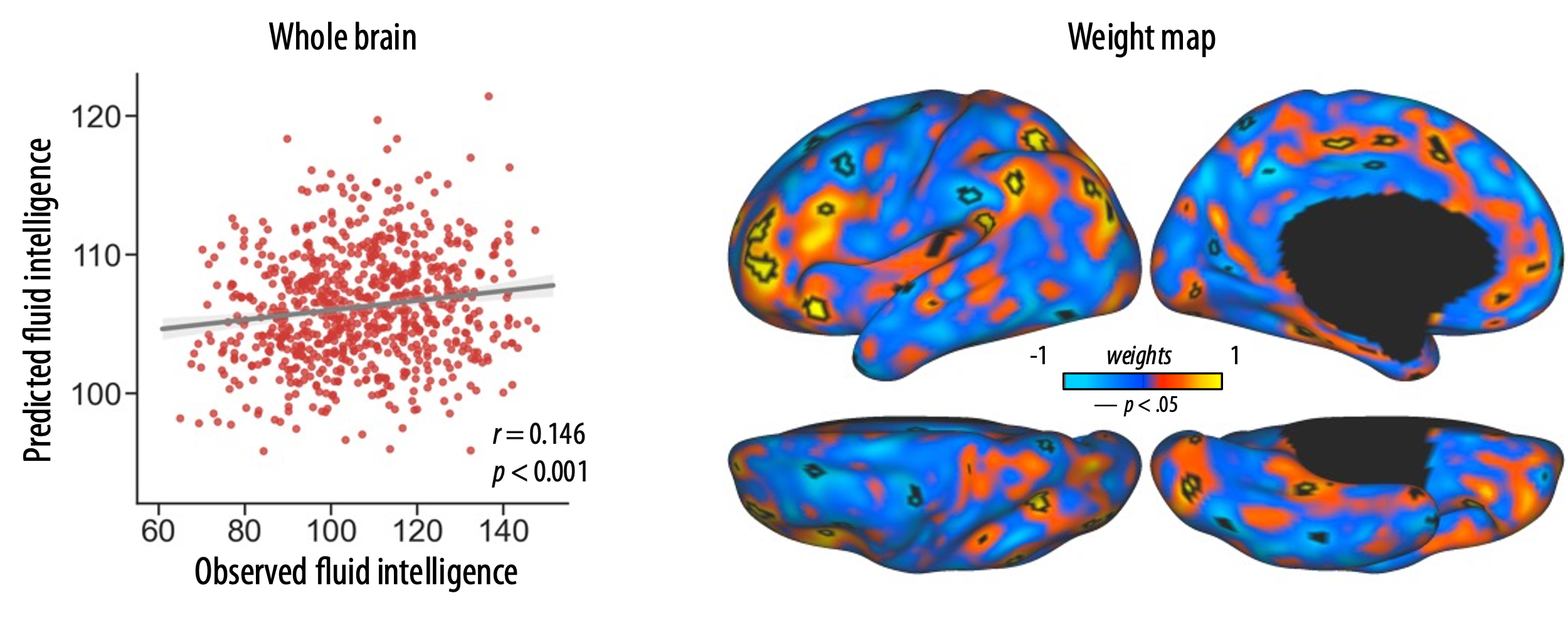

Fig. S10. The prediction of fluid intelligence by using leave-one-subject-out cross-validation. We applied support vector regression (SVR) with leave-one-subject-out cross-validation to test the generalization of vertexwise association between between-hemisphere functional distance and fluid intelligence. The Pearson correlation between predicted and original values reached 0.146. A bootstrap method by randomizing the original labels 5,000 times to reveal the brain regions that made significant contribution to the model. The regions with uncorrected *p* < 0.05 were shown with black contours.

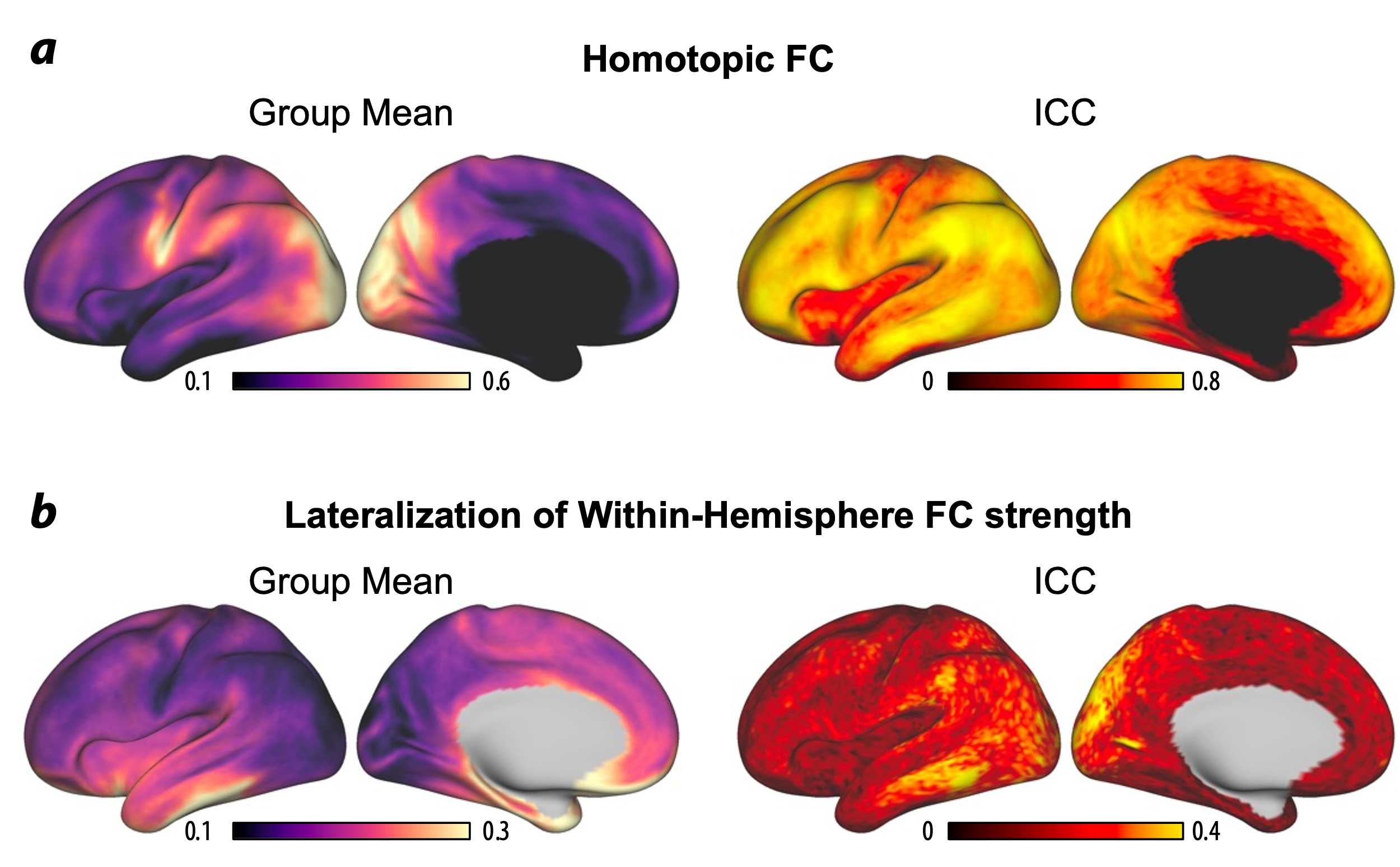

Fig. S11. The spatial patterns of other functional measures for hemispheric differences.

|  | Main (n = 755) | | Validation (n = 755) | |
| --- | --- | --- | --- | --- |
|  | *r* | *p* | *r* | *p* |
| Fluid composite | 0.125 | <0.001** | 0.118 | 0.001** |
| PicSeq | 0.088 | 0.016* | 0.084 | 0.022* |
| CardSort | 0.134 | <0.001** | 0.132 | <0.001** |
| Flanker | 0.065 | 0.076 | 0.056 | 0.124 |
| ProcSpeed | 0.078 | 0.032* | 0.063 | 0.084 |
| ListSort | 0.044 | 0.224 | 0.053 | 0.146 |

Table S1. Correlations with the fluid composite score and its subdomains

The associations between global between-hemisphere functional distance and cognitive scores were calculated by using the Pearson correlation coefficient, controlling for age, sex, handedness and head motion (mean framewise displacement).

Abbreviations: PicSeq = Picture sequence memory task for episodic memory; CardSort = Dimensional change card sort task for executive cognitive flexibility; ProcSpeed = Salthouse pattern comparison task for processing speed; ListSort = List sorting task for working memory.

*Uncorrected *p* < 0.05; ** Bonferroni corrected *p* < 0.05

|  | Main (n = 755) | | Validation (n = 755) | |
| --- | --- | --- | --- | --- |
|  | *r* | *p* | *r* | *p* |
| Age | 0.159 | <0.001** | 0.138 | <0.001** |
| Handedness | 0.052 | 0.155 | 0.057 | 0.119 |
| Total brain volume | 0.238 | <0.001** | 0.260 | <0.001** |
|  | *F*_(1,749)_ | *p* | *F*_(1,749)_ | *p* |
| Sex | 5.55 | 0.019* | 7.54 | 0.006* |

Table S2. The influences of demographics and brain size on global between-hemisphere functional distance.

*Uncorrected *p* < 0.05; ** Uncorrected *p* < 0.001

|  | Homotopic FC (n = 751) | | Lateralization of within-hemisphere integration (n = 751) | |
| --- | --- | --- | --- | --- |
|  | *r* | *p* | *r* | *p* |
| Total brain volume | 0.115 | 0.002** | 0.034 | 0.349 |
| Fluid Composite | -0.020 | 0.578 | 0.012 | 0.746 |
| PicSeq | 0.039 | 0.293 | 0.080 | 0.030* |
| CardSort | -0.071 | 0.054 | -0.032 | 0.376 |
| Flanker | 0.008 | 0.824 | 0.005 | 0.882 |
| ProcSpeed | -0.088 | 0.016* | -0.014 | 0.696 |
| ListSort | 0.050 | 0.171 | 0.008 | 0.831 |

Table S3. Correlation analyses for other types of functional lateralization.

The associations between global between-hemisphere functional distance and cognitive scores were calculated by using the Pearson correlation coefficient, controlling for age, sex, handedness and head motion (mean framewise displacement).

Abbreviations: PicSeq = Picture sequence memory task for episodic memory; CardSort = Dimensional change card sorting task for executive cognitive flexibility; ProcSpeed = Salthouse pattern comparison task for processing speed; ListSort = List sorting task for working memory.

*Uncorrected *p* < 0.05; ** Bonferroni corrected *p* < 0.05

| Projected to **left** hemisphere | | |  | | Projected to **right** hemisphere | | | |
| --- | --- | --- | --- | --- | --- | --- | --- | --- |
| MMP-ID | MMP-Name | Number | |  | | MMP-ID | MMP-Name | Number |
| 149 | PFm | 44 | |  | | 149 | PFm | 38 |
| 85 | a9-46v | 24 | |  | | 85 | a9-46v | 24 |
| 171 | p47r | 14 | |  | | 25 | PSL | 18 |
| 40 | 24dd | 14 | |  | | 111 | AVI | 12 |
| 111 | AVI | 11 | |  | | 40 | 24dd | 12 |
| 84 | 46 | 7 | |  | | 84 | 46 | 11 |
| 95 | OFC | 7 | |  | | 117 | AIP | 8 |
| 25 | PSL | 6 | |  | | 82 | IFSa | 7 |
| 150 | PF | 6 | |  | | 37 | 5mv | 6 |
| 102 | OP2-3 | 6 | |  | | 9 | 3b | 5 |
| 9 | 3b | 5 | |  | | 102 | OP2-3 | 5 |
| 127 | PHA3 | 5 | |  | | 171 | p47r | 5 |
| 6 | V4 | 5 | |  | | 127 | PHA3 | 5 |
|  |  |  | |  | | 81 | IFSp | 5 |

Table S4. Brain locations of significant vertices.

To identify the attribution of the significant vertices related to fluid intelligence, we projected them to the left and right hemispheres. The regions were located by using the multimodal parcellation (MMP) and are listed in decreasing order with their significant numbers included. Regions with fewer than 5 vertices were excluded.
